# Flight muscle allocation diverges between two moth families with distinct flight strategies

**DOI:** 10.64898/2026.08.24.746726

**Authors:** Joanna Baker, Ethan Wold, Leo Wood, Brett Aiello, Simon Sponberg

## Abstract

An animal’s musculature must support its specific biomechanical needs, so muscle morphology and volume allocation may adapt when locomotor strategies diversify. We examined muscle size and morphology in two sister families of bombycoid moths, wild silkmoths (*Saturniidae*) and hawkmoths (*Sphingidae*), that have diverged in wingbeat frequency, wing morphology, and behavior. Although both families rely on the same muscles to power and steer flight, they may distribute muscle volume differently to prioritize distinct functions. We hypothesized that flight power muscle proportions are larger in hawkmoths and increase with wingbeat frequency, helping meet inertial power demands of high-frequency maneuverable flight. We also hypothesized that some individual muscles diverge in proportional volume and area to support distinct wing control strategies. To test our hypotheses, we took µCT scans of twenty bombycoid species and quantified volumes and geometries of six flight muscle pairs. As expected, flight power muscle proportions positively correlate with wingbeat frequency and are generally greater in hawkmoths. Two of three steering muscles diverge substantially in relative volume and area between families. Most muscles exhibit greater length in silkmoths and greater cross-sectional area in hawkmoths. Finally, the dorsal oblique (DO) muscle diverges exceptionally in size and morphology, being highly developed in hawkmoths and smaller or absent in silkmoths. This unexpected difference supports the DO having an underappreciated role in flight control, possibly via shaping indirect strain propagation in the elastic thorax. We show that muscle volume distribution parallels bombycoids’ divergent flight strategies, demonstrating how muscle allocation can adapt for specialized functional goals.

## 1 Introduction

Animal movement involves precise coordination of many parts including limbs, skeletal structures, muscles, and sensorimotor circuits. Determining how these integrated components evolve in conjunction to alter whole-body motion is key to understanding patterns in locomotor evolution (Farina et al., 2019; Wainwright & Price, 2016). Flying insects exhibit extraordinary variation in anatomy and behavior, which likely entailed evolutionary changes to the thoracic flight muscles which drive the wingbeats (Dudley, 2000; Marden, 2000). Flight muscle anatomy and function are well-studied in some model insects (Dickinson & Tu, 1997; Eaton, 1988; Kammer, 1967, 1970, 1971; Ortega et al., 2019), and the relative sizes of flight muscles can vary substantially across insect orders (Pringle, 1968). However, to understand how the relative size and arrangement of flight muscles adapts to achieve specialized functional goals, we must harness comparative study of insects which evolved distinct flight strategies while sharing the same underlying muscular organization. Bombycoid moths provide a compelling model clade for this purpose (Aiello et al., 2026), presenting an opportunity to investigate how flight muscle allocation adapted during a specific evolutionary divergence in flight morphology and behavior.

The superfamily Bombycoidea contains two sister families — wild silkmoths (*Saturniidae*; hereafter silkmoths) and hawkmoths (*Sphingidae*) — which evolved distinct flight strategies [Fig. 1A]. Silkmoths, with larger, lower aspect ratio (AR) wings and typically slower wingbeat frequencies ( 5-25 Hz), perform erratic pitching and bobbing which likely aids in predator avoidance (Aiello, Sikandar, et al., 2021; Aiello, Tan, et al., 2021; Humphries & Driver, 1970). Silkmoths feed only as caterpillars and have finite energy stores as winged adults, despite flying significant distances to locate mates and host plants (Janzen, 1984; Tuskes et al., 1996). Conversely, hawkmoths have smaller, higher AR wings and perform controlled, agile flight at higher frequencies (~20-70 Hz) (Aiello, Sikandar, et al., 2021; Aiello, Tan, et al., 2021). Fast flapping facilitates rapid and precise maneuvers, allowing many hawkmoths to hover-feed similarly to hummingbirds (Janzen, 1984; Willmott & Ellington, 1997). However, hawkmoths sacrifice aerodynamic effiiciency to maintain maneuverability at higher frequencies (Gau et al., 2021, 2022; Wold, Aiello, et al., 2024). Because silkmoths and hawkmoths differ in morphology and behavior while spanning an order-of-magnitude difference in wingbeat frequency, they may experience different demands on their thoracic flight musculature (Marden, 2000). We aim to understand how flight muscles differ in size and morphology between these families to support their contrasting flight strategies. Although *Lepidoptera* possess dozens of thoracic muscles (Eaton, 1988), a few large muscle pairs in the mesothorax comprise the basic mechanism for producing flight power and wingstroke-to-wingstroke control [Fig. 1B-C, Table 1] (Ortega et al., 2019). Indirect flight power muscles — the dorsolongitudinal (DLM) and dorsoventral (DVM) — generate the downstroke and upstroke by indirectly deforming the thoracic exoskeleton, while direct steering muscles — the basalar, subalar, and third axillary — act directly upon wing hinge sclerites to position the wings (Chapman, 1998; Dickinson & Tu, 1997; Kammer, 1967). Additional direct muscles that have functional roles in other insects (e.g. Lindsay et al. (2017)) tend in *Lepidoptera* to share motor neuron innervation with the three main direct steering muscles, or appear reduced and of tonic fiber types, suggesting comparatively subtle contributions to wing control (Eaton, 1988; Kammer, 1971; Kammer & Rheuben, 1981; Ortega et al., 2019). Accordingly, the activity of the five main muscle pairs alone allows near-complete reconstruction of flight kinematics in the hawkmoth *Manduca sexta* (Ortega et al., 2019, 2023; Yang et al., 2022). Still, some other muscles are thought to play important and underappreciated roles in flight, particularly the dorsal oblique (DO), another indirect muscle which is prominent in *M. sexta* though much smaller than the DLM and DVM [Fig. 1B]. It has an unresolved function but may assist in control (Kammer, 1967, 1971; Ortega et al., 2019; Pringle, 1968). Together, these six muscles provide a compact and comprehensive set to examine for flight adaptations (Ortega et al., 2019; Pringle, 1968).

**Figure 1.**
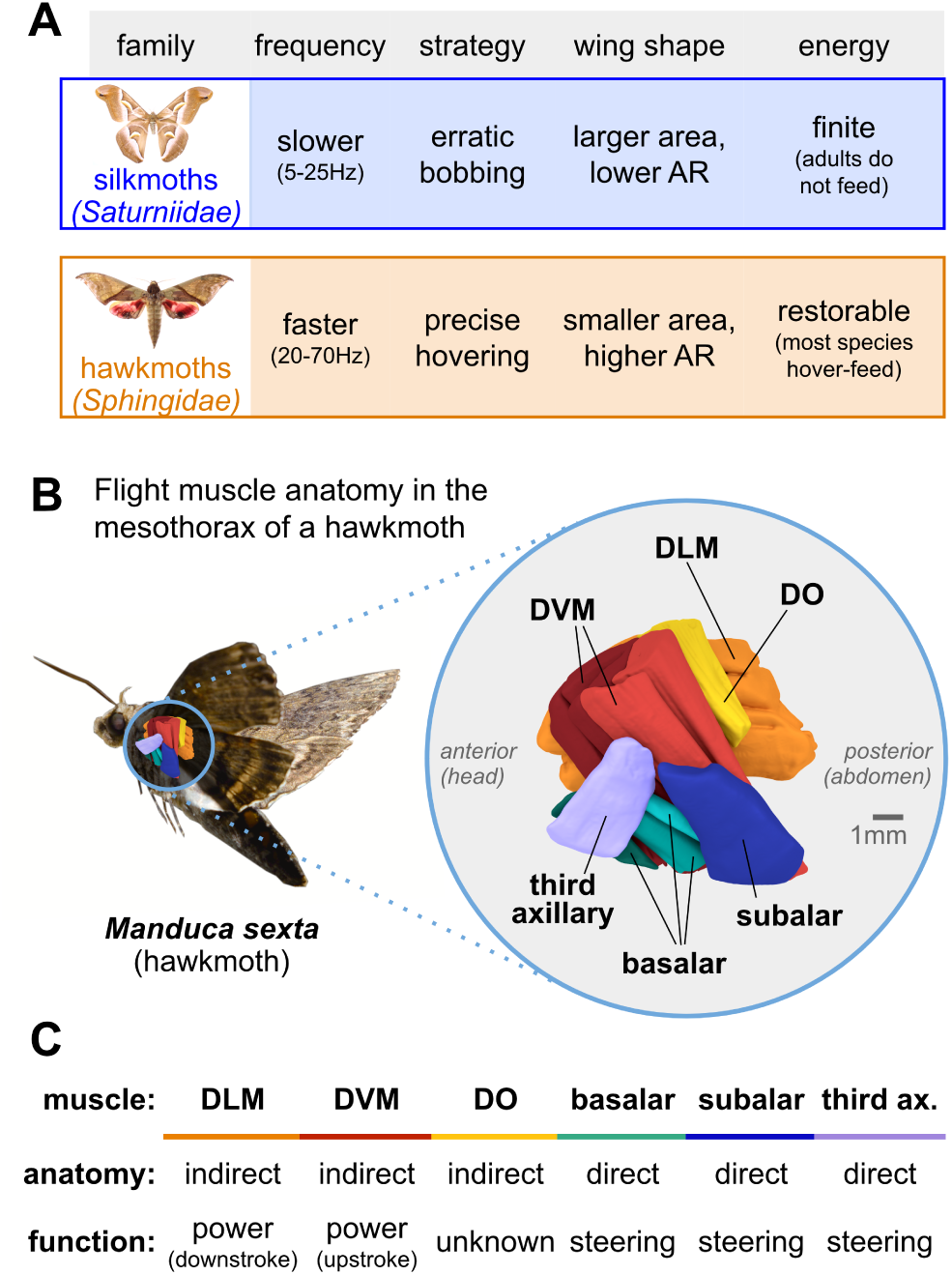
Silkmoths and hawkmoths may allocate their flight muscle differently. **A.** Wild silkmoths (*Saturniidae*) and hawkmoths (*Sphingidae*) are closely-related moth families which have diverfed in wingbeat frequency, flight strategy, wing shape, and energy budget. In this study, we examine whether flight muscle proportions parallel these known differences. **B.** We used µCT to image moth thoraces and segment six flight muscle pairs. Mesothoracic muscle anatomy is shown for the model hawkmoth, *Manduca sexta*. Across figures, multicolor visualizations denote muscle identity as specified here; these are unrelated to family color assignments (blue for *Saturniidae*, orange for *Sphingidae*) used in family comparison visualizations. **C.** Flight muscles have either direct or indirect anatomy, depending on whether or not they attach to the wing hinge. Indirect power muscles (DLM and DVM) do not attach to the wing hinge and instead deform the exoskeleton to power flight, while direct steering muscles (basalar, subalar, and third axillary) control wing positioning through the wing hinge. The dorsal oblique (DO) muscle has indirect anatomy but unknown function.

**Table 1.**
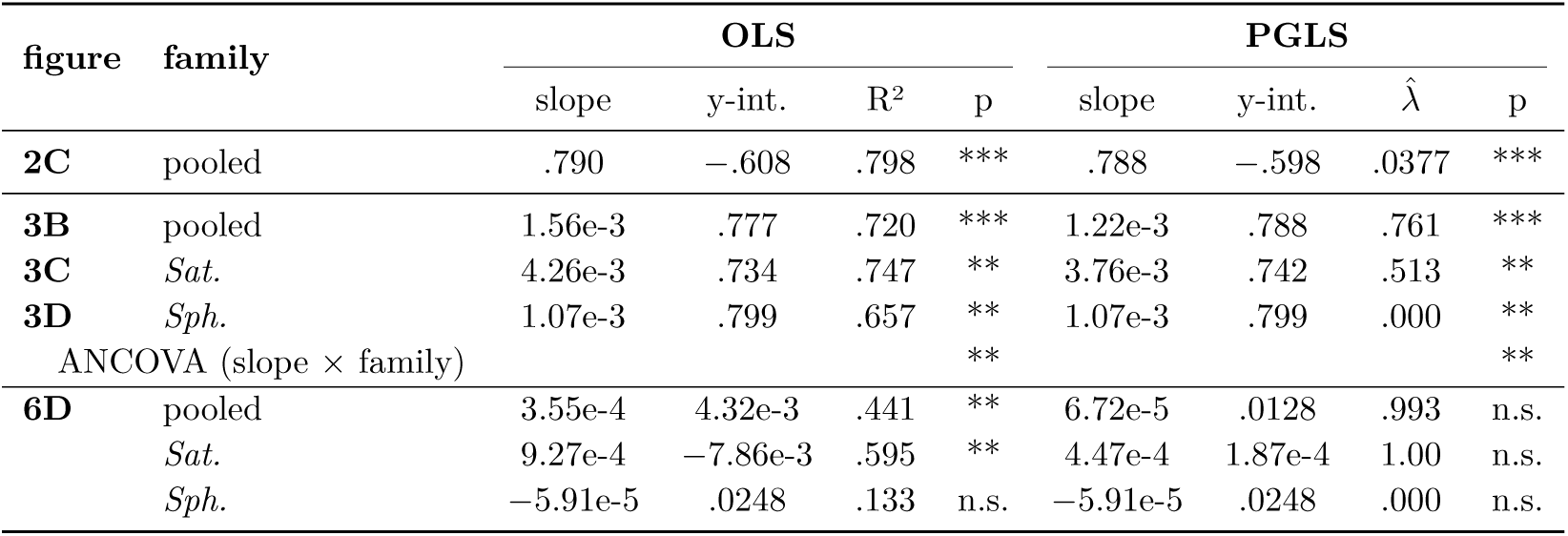
Regression statistics for. **Figures 2, 3, and 6.** All regressions use ordinary least squares (OLS) and phylogenetic generalized least squares (PGLS). *λ*^^^ denotes the maximum-likelihood estimate of Pagel’s *λ* fitted as part of each PGLS model, reflecting phylogenetic covariance remaining in model residuals after accounting for predictors. **Fig. 2C**: log total flight muscle volume (mm^3^) vs. log body mass (mg); *n* = 20; slope represents nondimensional log-log scaling exponent; isometry predicts scaling exponent = 1. **Fig. 3**: indirect muscle proportional volume vs. wingbeat frequency (Hz); 3B = all species combined, 3C = *Saturniidae*, 3D = *Sphingidae* (*n* = 10 per family). Slope units in Hz*^−^*^1^. ANCOVA row reports the significance of the wingbeat frequency family interaction term, used to evaluate whether regression slopes differ between *Saturniidae* and *Sphingidae*. **Fig. 6D**: dorsal oblique proportional volume vs. wingbeat frequency (*n* = 10 per family). Slope units in Hz*^−^*^1^. Significance: *** p < 0.001, ** p < 0.01, * p < 0.05, n.s. p ≥ 0.05.

Muscle volume, length, and cross-sectional area help measure physical constraints on muscle capacity. Flying insects face strong mechanical and physiological pressures to consolidate muscle size, because muscles are the primary energy-consuming tissues during flight and are costly to use, maintain, and carry (Marden, 2000). This makes volume an informative metric for quantifying how resources and space are invested into each muscle and allocated across the flight musculature. Greater volume broadly corresponds to more contractile tissue, and isometric scaling predicts that potential mechanical power output increases proportionally with muscle volume (Woittiez et al., 1984). This is because volume is the product of cross-sectional area, which scales with force production ability, and fiber length, which scales with contraction velocity (Hill, 1938; Josephson, 1999; Pennycuick & Rezende, 1984). Therefore, decomposing volume into its area and length components can provide insights into how force production capacity and velocity generation capacity drive overall investment into each muscle (Lieber & Ward, 2011; Orsbon et al., 2018; Woittiez et al., 1984). Morphology alone cannot predict *in vivo* muscle performance, which depends on numerous factors including fiber composition and architecture, sarcomere physiology, and neural activation patterns (Ellington, 1985; Roberts et al., 2019; Rome & Lindstedt, 1998; Sponberg & Daniel, 2012). Nevertheless, comparing muscle size and morphology between silkmoths and hawkmoths may uncover distinct approaches towards balancing resource investment across flight functions.

We examined how flight muscle allocation varies with flight strategy across twenty silk- moth and hawkmoth species. Using µCT reconstruction, we quantified the volumes and cross-sectional areas of the major muscles that drive the wingbeat. Because these six flight muscles play distinct roles in power production and wing control, we expected that the contrasting flight strategies of silkmoths and hawkmoths would be accompanied by corresponding differences in muscle size and morphology. Our study tested three specific hypotheses:

1. The combined volume proportion of the indirect flight muscles is greater in hawkmoths and increases with wingbeat frequency, reflecting the heightened power demands of high-frequency controlled hovering.
2. For each of the three direct steering muscles, silkmoths and hawkmoths show significant differences in average volume proportions and normalized cross-sectional areas, given the families’ extensive differences in wing steering kinematics.
3. The dorsal oblique muscle occupies a larger volume and area in hawkmoths, consistent with the muscle’s possible role in flight control. These hypothesized outcomes would support the idea that flight strategy evolution involves non-uniform shifts in muscle allocation, with changes concentrated in muscles that experience divergent functional demands.

## 2 Methods

### 2.1 Specimens

We used µCT to image the mesothoracic musculature of twenty bombycoid species. We examined ten wild silkmoths (*Saturniidae*) and ten hawkmoths (*Sphingidae*), sampling broadly within each family. Species from the silkmoth family include: *Actias luna* (AL), *Antheraea polyphemus* (AP), *Automeris io* (AI), *Callosamia angulifera* (CA), *Citheronia regalis* (CR), *Eacles imperialis* (EI), *Hemileuca maia* (HM), *Hyalophora euryalus* (HE), *Samia cynthia* (SC), and *Saturnia walterorum* (SW). Species from the hawkmoth family include: *Amorpha juglandis* (AJ), *Amphion floridensis* (AF), *Eumorpha achemon* (EA), *Hemaris thysbe* (HT), *Hyles lineata* (HL), *Manduca sexta* (MS), *Paonias myops* (PM), *Proserpinus terlooi* (PT), *Smerinthus ophthalmica* (SO), and *Sphinx chersis* (SpC).

We acquired a single specimen of each species, sourcing specimens live from breeders or preserved in ethanol from laboratory collections. Specimens acquired live were anesthetized via cold exposure. After weighing each insect, we isolated its thorax by detaching the head, wings, abdomen, and legs. Using a thin needle, we punctured shallow holes in the sides of the thorax to enable stain permeation. Before staining, we submerged each thorax in ethanol for at least 24 hours. Then, we soaked each thorax in an aqueous solution of 10% phosphotungstic acid (PTA), a contrast-enhancing stain (O et al., 2019; Swart et al., 2016), for three weeks prior to scanning.

We only scanned specimens without external damage to the thorax. We also examined each scan before complete segmentation and analysis to confirm that muscle morphology was intact and symmetrical. Two additional species were scanned but the specimens were excluded from further analysis upon observing abnormal tissue deterioration and asymmetry. These quality-control steps ensured consistent tissue integrity in all analyzed specimens.

We did not record individuals’ sex, as many preserved specimens could not be sexed. Although sexual dimorphism does exist among *Bombycoidea*, much of sex-based variation is sensory and behavioral; biomechanically-relevant differences are limited by the constraint that both sexes fly (Agosta & Janzen, 2005; Camargo et al., 2015; Janzen, 1984). Furthermore, because we focus on relative muscle allocation rather than absolute muscle size, our metrics of interest are presumably independent of body size differences between sexes.

### 2.1.1 µCT

We imaged each prepared thorax using a SCANCO µCT 50 machine. Specimens were scanned at 70 kV, 115 µA, and 500 ms, with a 0.1-µm aluminum filter.

#### **2.1.2** Muscle selections

We focused our analysis on six bilateral muscle pairs per moth [Table 1], five of which almost entirely describe the wing motion of the hawkmoth *M. sexta* (Ortega et al., 2019, 2021, 2023; Yang et al., 2022). The mesothoracic dorsolongitudinal muscle (DLM; Eaton (1988): II dl1) and mesothoracic dorsoventral muscle (DVM; II dv1-5) indirectly power the downstroke and upstroke respectively, while the basalar (II pv1-3), subalar (II pv4-5), and third axillary (II pd2a-b) perform direct steering functions (Chapman, 1998; Dickinson & Tu, 1997; Kammer, 1967, 1971). A sixth muscle, the DO (II dl2), is not as well-studied, but is prominent in the *M. sexta* mesothorax and might be involved in flight control (Eaton, 1988; Kammer, 1967, 1971; Ortega et al., 2019; Pringle, 1968). Fig. 1B-C shows the anatomy and broad functional classifications of these muscles.

Because muscles do not act in isolation, their functions depend heavily on coordinating their activation timing with other muscles (Dickinson & Tu, 1997; Ortega et al., 2019; Wood et al., 2024; Yang et al., 2022). Therefore, each flight muscle has complex, multifaceted roles which may go beyond basic categorization as power or steering (Chapman, 1998; Kammer, 1971; Sponberg & Daniel, 2012; Springthorpe et al., 2012; Wood et al., 2024). In interpreting the implications of muscle size variation, we focus mainly on anatomical groups and primary established functions, but discuss other known or speculated roles for muscles when relevant.

#### **2.1.3** Digital segmentation

To perform digital segmentation of these twelve muscles from the µCT scans, we used the open-source image computing platform 3D Slicer (5.6.2) (Fedorov et al., 2012) with the Slicer Extra Effects package (Lasso, 2025) to capture the structure of the flight muscles and calculate their volumes. Muscles with multiple subunits were considered as one muscle in analyses. We used manual segmentation to cleanly isolate muscle geometries while preserving fine structural features. See Supplementary Information SI.1 for detailed methods. Following segmentation, we exported object files to Blender for rendering.

### **2.2** Analysis

#### **2.2.1** Phylogeny

We assembled a genus-level composite time-resolved phylogeny covering all taxa in our study [Fig. 2A] using the R packages *phytools* (2.3.0) and *ape* (5.8). The tree is resolved to the genus level because the source trees contained some taxa with the same genus as a study specimen but a different species. The phylogeny from Aiello, Tan, et al. (2021) served as our base tree, as it contains nearly all genera we sampled. For the two hawkmoth genera not represented in that phylogeny, *Amorpha* and *Proserpinus*, we obtained tree position and branch length data from Kawahara and Barber (2015). Using the *bind.tip* function, we grafted these nodes onto the pruned tree while maintaining time calibration.

**Figure 2.**
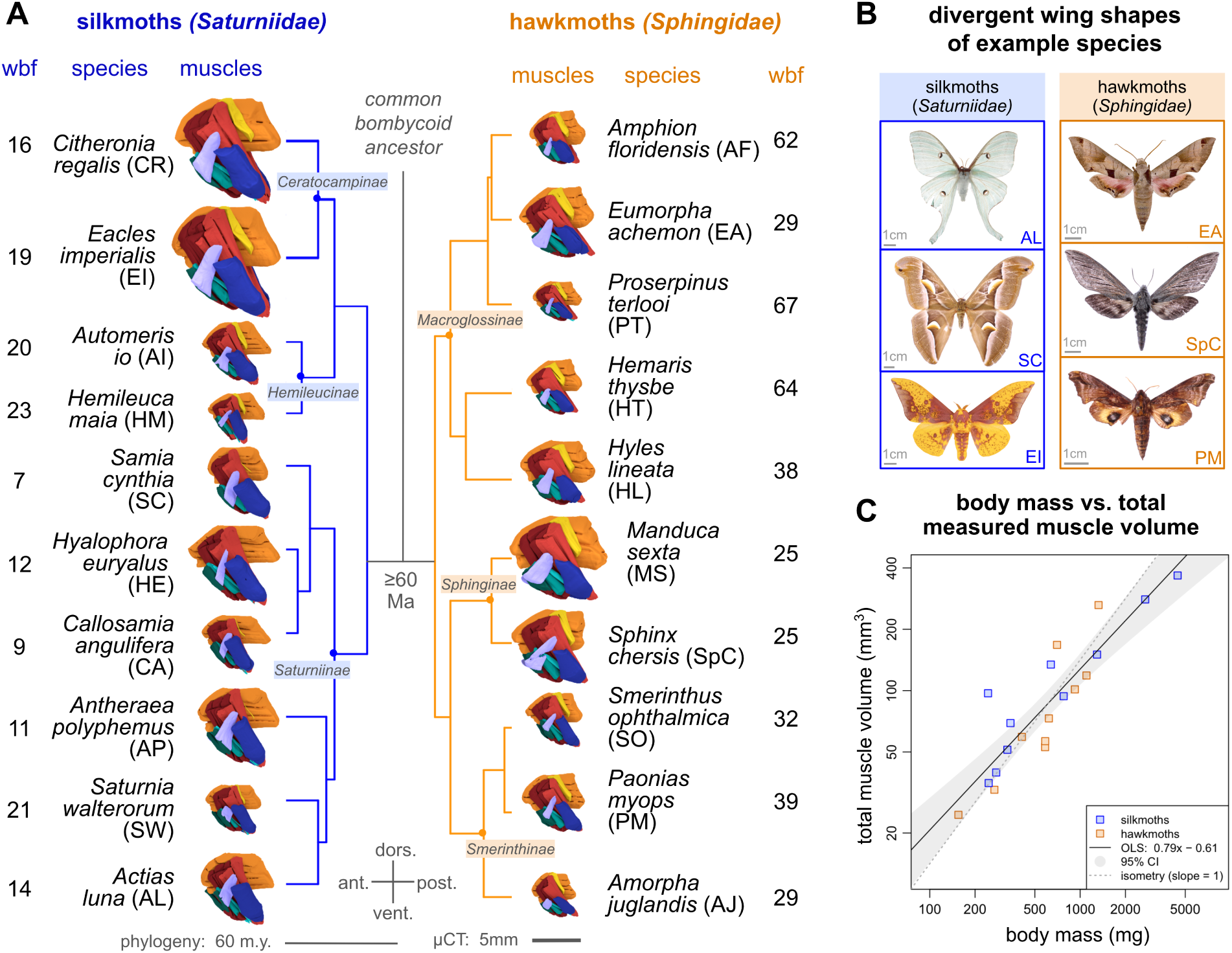
µCT imaging reveals the thoracic flight muscle anatomy of twenty bombycoid moths. **A.** We scanned ten silkmoths (blue, left) and ten hawkmoths (orange, right), sampling broadly within each family. Renderings of six flight muscles, to scale, are presented alongside a time-resolved phylogeny, with species names, abbreviations, and wingbeat frequency (wbf) estimates in Hz. Subfamilies are labeled at nodes. **B.** Silkmoths and hawkmoths possess divergent wing morphologies (Aiello, Tan, et al., 2021); representative species are visualized with 1cm scale bars. **C.** Total measured muscle volume correlates positively with body mass across bombycoids on a log-log scale [Table 2], validating total muscle volume as a proxy for normalizing individual muscle volumes to body size. An F-test reveals that the scaling exponent of this regression, 0.79 (p < 0.05 for OLS and PGLS), is hypometric relative to the expectation of isometry (scaling exponent = 1).

**Table 2.** Flight muscle volume proportions in each family. Mean flight muscle volume proportions and 95% confidence intervals for six muscle pairs among *Saturniidae* (silkmoths, n = 10) and *Sphingidae* (hawkmoths, n = 10). Proportion is calculated for each individual as bilateral muscle volume divided by total measured flight muscle volume. Indirect muscle proportion is the sum of the dorsolongitudinal (DLM), dorsoventral (DVM), and dorsal oblique (DO) muscle proportions. Direct muscle proportion is equivalent to 100% indirect. Welch’s t-tests compare family means while treating species as independent observations; phylogenetic ANOVA incorporates shared evolutionary history. Significance: *** p *<* 0.001, ** p *<* 0.01, * p *<* 0.05, n.s. = not significant. sectional area was detected only for the DVM [Supp. Fig. S2]. We discuss these scaling patterns further in Supplementary Information SI.2.

|  |  | *indirect* | DLM | DVM | DO | basalar | subalar | third ax. |
| --- | --- | --- | --- | --- | --- | --- | --- | --- |
| ***Sat.*** | mean (%) | 79.9 | 44.2 | 35.0 | 0.62 | 4.82 | 13.8 | 1.55 |
|  | 95% CI | [77.9–81.8] | [42.9–45.5] | [34.3–35.8] | [0.15–1.09] | [4.34–5.29] | [12.4–15.2] | [1.18–1.92] |
| ***Sph.*** | mean (%) | 84.3 | 50.4 | 31.7 | 2.23 | 4.76 | 6.97 | 3.99 |
|  | 95% CI | [82.7–85.9] | [47.6–53.2] | [29.6–33.8] | [2.04–2.43] | [4.46–5.05] | [6.09–7.85] | [3.46–4.52] |
| Welch’s t-test |  | *** | *** | ** | *** | n.s. | *** | *** |
| phy. ANOVA |  | n.s. | n.s. | n.s. | * | n.s. | * | * |

#### **2.2.2** Wingbeat frequency estimates

Free-flight wingbeat frequencies have been measured in some study taxa, but not all. For consistency across all species, we estimated wingbeat frequencies (*n*) using body mass (*m*) and wing area (*A*) data from Aiello, Tan, et al. (2021). Deakin (2010) provides two formulae for calculating *n* in Hz, given *m* in grams and *A* in mm^2^:

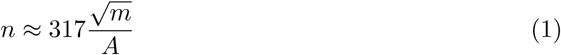

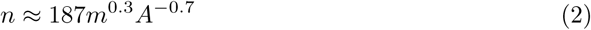

These two formulae yielded slightly different values, with the second formula returning a higher result in nearly all species. We opted for a middle-ground frequency estimate in Hz by taking the average of the two calculated values. To validate this approach, we compared estimated and actual frequencies in species with measured free-flight frequencies, referencing two prior studies (Aiello, Sikandar, et al., 2021; Wold, Aiello, et al., 2024). Measured and calculated frequencies had a mean difference of -0.85 Hz and a mean absolute difference of 1.87 Hz. Because these values lie within the range of frequency variation observed in individual bombycoid moths (Gau et al., 2021; Yu et al., 2020), this method suffiiciently approximates actual flight frequencies. Body mass, wing area, calculated frequencies, and measured frequencies are reported by species in Supplementary Table S1.

#### **2.2.3** Volume proportions and muscle geometry

We extracted the individual muscle volumes in mm^3^ from 3D Slicer via *Quantification Segment Statistics*. For each bilateral muscle pair, we summed the two volumes to obtain a single combined volume for each muscle [Supp. Table S2].

For a given muscle pair or group, we obtained its proportional volume [Supp. Table S3] — a nondimensional quantity — by dividing its bilateral volume over the total muscle volume in that specimen, the sum of the twelve measured muscle volumes. Total muscle volume correlates closely with body mass [Fig. 2C], making it an appropriate normalization for comparing muscle investment across individuals of different body sizes. Here, total muscle volume only includes the six pairs studied, and therefore does not reflect the animal’s true total muscle volume.

We used 3D Slicer to measure the length of each muscle along its longitudinal axis (Eaton, 1988; Tu & Daniel, 2004). For the DLM, DVM, basalar, and subalar, we measured subunits separately and averaged the lengths. We then estimated cross-sectional areas by dividing unilateral muscle volume by muscle length [Supp. Tables S4 & S5]. These estimates may not precisely reflect physiological cross-sectional area, as we did not measure fiber pennation angle (Martin et al., 2020; Roberts et al., 2019). While our cross-sectional area measurements are thus not suitable for exact calculations of muscle force capacity, they provide a reasonable approximation for comparing muscle areas among species.

#### **2.2.4** Statistical methods

We analyzed data in R (4.4.1) with RStudio using base statistical functions and phylogenetic comparative methods. We performed Ordinary Least Squares (OLS) linear regressions to examine the scaling of body mass with total muscle volume across all 20 species [Fig. 2C, Table 1]. We also used OLS regression to examine relationships between wingbeat frequency and indirect muscle proportion across all species [Fig. 3B] and within each family [Fig. 3C-D, Table 1], as well as between wingbeat frequency and dorsal oblique muscle proportion across all species and within each family [Fig. 6D, Table 1]. Regression coeffiicients, coeffiicients of determination (*R*_2_), and 95% confidence intervals were calculated for all models [Table 1]. We used phytools (2.3.0), ape (5.8), and caper (1.0.3) to conduct phylogenetic comparative analyses. Accompanying all OLS regressions, we performed Phylogenetic Generalized Least Squares (PGLS) regressions [Table 1] to determine whether relationships remained significant after accounting for phylogenetic non-independence among species (Grafen, 1989; Symonds & Blomberg, 2014). For each PGLS model, Pagel’s *λ*^^^ was estimated by maximum likelihood; this parameter describes the degree of phylogenetic covariance remaining in model residuals after accounting for predictor variables (Pearse et al., 2023).

**Figure 3.**
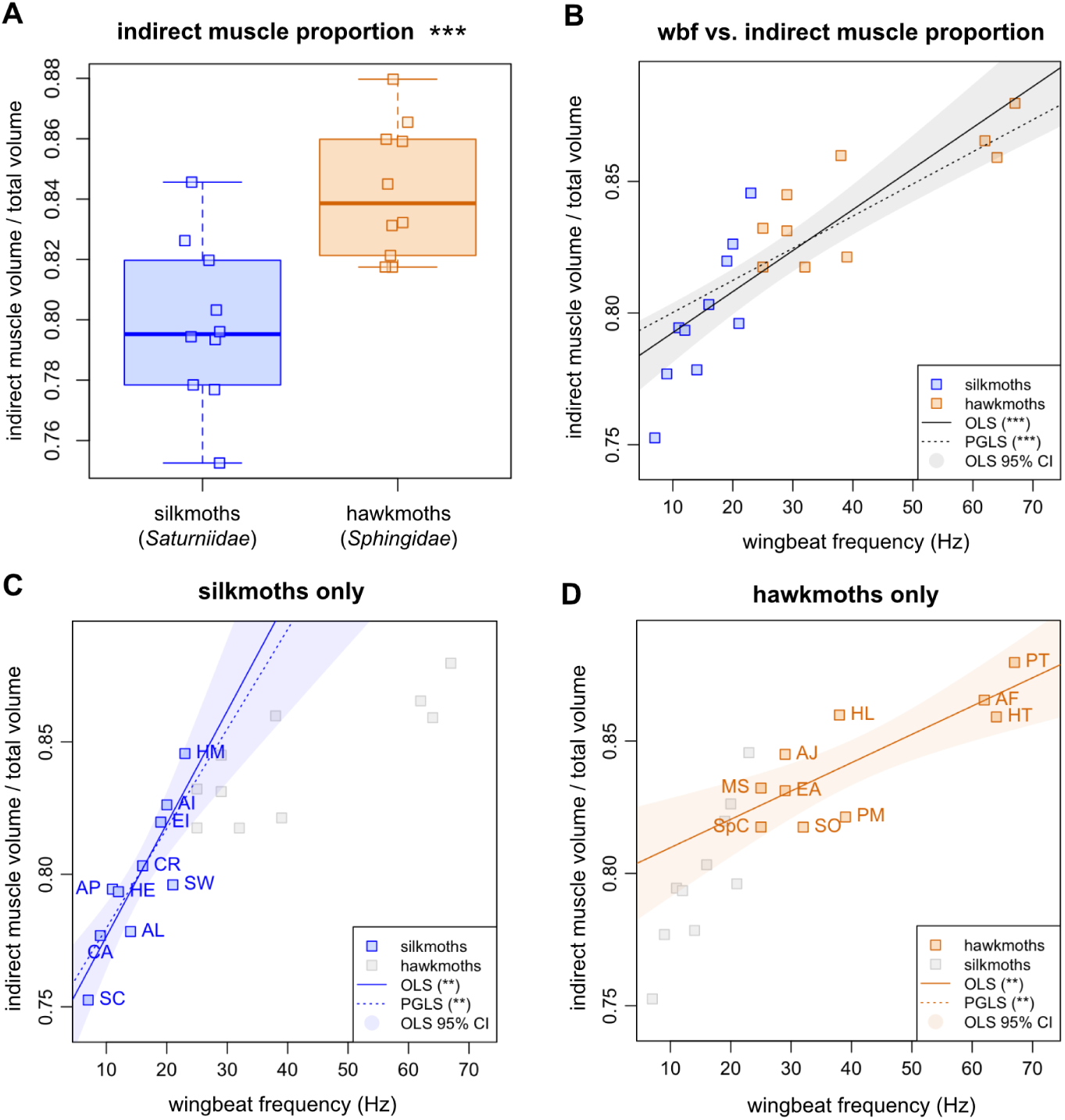
Faster-flapping bombycoid moths have larger proportions of indirect flight muscles which power flapping. **A.** Hawkmoths tend to have larger proportions of indirect flight muscle volume than silkmoths; inversely, silkmoths have larger proportions of direct flight muscle volume (t-test p < 0.001). However, this divergence is not significant under phylogenetic ANOVA [Table 2]. **B.** Wingbeat frequency (wbf) positively correlates with indirect muscle proportion (slope = 0.0016 Hz^−1^), including after phylogenetic correction (slope = 0.0012 Hz^−1^), suggesting that higher-frequency fliers allocate more muscle capacity towards flight power [Table 1]. **C.** The frequency-indirect proportion relationship remains significant when examining silkmoths only and has a steeper slope (OLS 0.0043 Hz^−1^, PGLS 0.0038 Hz^−1^) than the pooled bombycoid relationship. Phylogenetic correction moderately reduces the slope (*λ*^^^ = 0.513), indicating that part of the relationship reflects shared evolutionary history within *Saturniidae*. **D.** The frequency-indirect proportion relationship also persists within hawkmoths only, but with a shallower slope, which does not change with phylogenetic correction (slope = 0.0011 Hz^−1^ for OLS & PGLS). The slopes differ significantly between families (ANCOVA interaction p < 0.01 for OLS and PGLS). Significance: *** p *<* 0.001, ** p *<* 0.01, * p *<* 0.05, n.s. = not significant.

To test whether frequency-proportion relationships exhibited different slopes in silkmoths and hawkmoths, we fit ANCOVA models of volume proportion as a function of wingbeat frequency, clade identity, and their interaction (volume proportion frequency clade) [Table 1]. Significance of the interaction term was evaluated with an F-test and used to assess the null hypothesis of equal slopes between clades. We implemented a phylogenetically informed equivalent using PGLS models that included wingbeat frequency, clade identity, and their interaction term. Significance of the interaction coeffiicient was likewise used to evaluate differences in slope between families.

To evaluate scaling allometry, we tested whether observed scaling exponents differed from isometric expectations using F-tests. Under geometric similarity, the expected scaling exponent for total muscle volume versus body mass is 1 (Dudley, 2000). Equivalent tests were performed for supplementary analyses of individual muscle volume versus body mass (isometric expectation = 1) and muscle cross-sectional area versus body mass (isometric expectation = 2/3) [Supp. Figs. S1 & S2, Supp. Table S6].

We performed two-tailed Welch’s t-tests to compare muscle volume proportions between silkmoths and hawkmoths [Table 2], analyzing indirect muscles as a group [Fig. 3A] followed by each muscle individually [Fig. 4]. For each family, we calculated mean volume proportions and corresponding 95% confidence intervals [Table 2]. Additionally, to accompany boxplots, we calculated median volume proportions and interquartile ranges (IQRs) within each family [Supp. Table S7].

**Figure 4.**
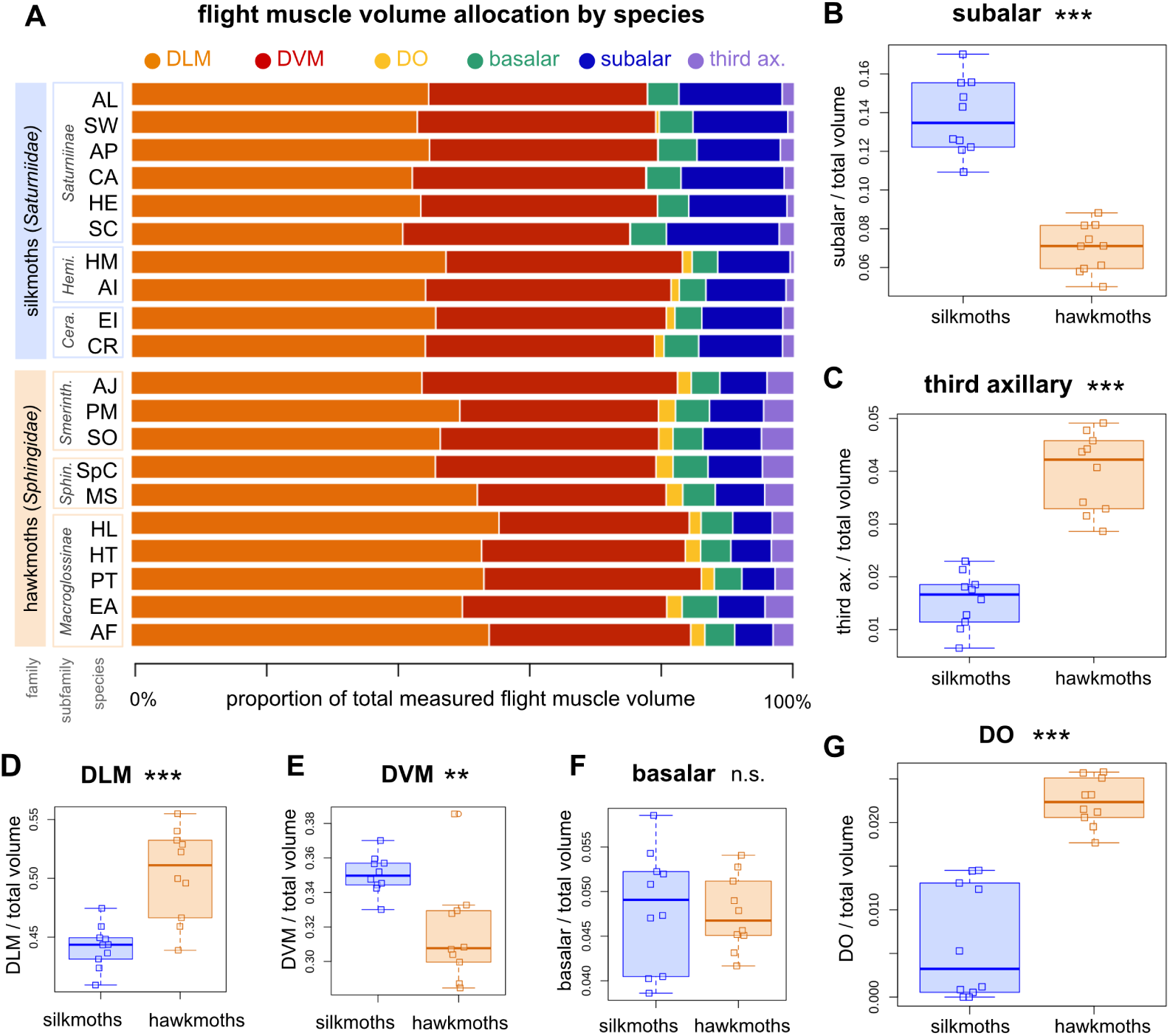
Volume allocation of individual flight muscles differs between silkmoths and hawkmoths. **A.** Volume proportions of six flight muscle pairs across twenty bombycoid species. Volume proportion is calculated as bilateral muscle volume divided by total muscle volume measured in that individual. **B–G.** Boxplots compare volume proportions between families for the subalar, third axillary, dorsolongitudinal (DLM), dorsoventral (DVM), basalar, and dorsal oblique (DO) muscles. We evaluated the significance of family divergences using Welch’s t-tests and phylogenetic ANOVA [Table 2]. Only the subalar, third axillary, and DO remain significant (p < 0.05) after phylogenetic correction. Muscle proportion values are reported by species in Supplementary Table S3. Asterisks indicate t-test significance: *** p < 0.001, ** p < 0.01, * p < 0.05, n.s. p ≥ 0.05.

To account for phylogenetic non-independence in family comparisons, we additionally used phytools to perform phylogenetic ANOVA using our genus-level pruned phylogeny [Table 2]. This was used to evaluate whether family differences in muscle proportion remained significant after accounting for shared evolutionary history (Garland et al., 1993).

To aid interpretation of phylogenetically-corrected statistical analyses, we quantified phylogenetic signal of muscle proportion data using Pagel’s *λ* and Blomberg’s K (Revell, 2013). *λ* and K quantify the degree to which variation in a trait is predicted by phylogenetic relatedness. Phylogenetic signal was evaluated for each muscle across the pruned phylogeny comprising both families, and within the subtree for each family [Table 4]. Pagel’s *λ* is distinct from the earlier metric, Pagel’s *λ*^^^, which describes residual phylogenetic covariance within a PGLS regression model (Pearse et al., 2023).

**Table 3.** Family centroids and multivariate comparisons of normalized muscle geometry. Values shown are family centroids for normalized muscle length (n.d. length) and normalized cross-sectional area (n.d. area) after scaling by total muscle volume to the 1/3 and 2/3 exponents, respectively. Hotelling’s two-sample T^2^ tests evaluate differences in joint length-area geometry between *Sphingidae* and *Saturniidae* [Fig. 5]. Larger T^2^ values indicate greater separation between families in normalized length-area morphospace and therefore stronger divergence in overall muscle geometry. Significance represents p-values adjusted using Holm correction for multiple comparisons across the six muscles. Corresponding univariate Welch’s t-tests for normalized length and area are reported in Supplementary Table 8. Significance: *** p < 0.001, ** p < 0.01, * p < 0.05, n.s. p ≥ 0.05.

|  |  | DLM | DVM | DO | basalar | subalar | third ax. |
| --- | --- | --- | --- | --- | --- | --- | --- |
| **n.d. length** | *Sat.* centroid | 1.410 | 1.479 | 0.505 | 0.836 | 1.118 | 0.633 |
|  | *Sph.* centroid | 1.221 | 1.377 | 0.651 | 0.722 | 0.916 | 0.533 |
| **n.d. area** | *Sat.* centroid | 0.157 | 0.150 | 0.005 | 0.029 | 0.061 | 0.012 |
|  | *Sph.* centroid | 0.207 | 0.184 | 0.017 | 0.033 | 0.038 | 0.037 |
| **Hotelling** | T<sup>2</sup> | 61.54 | 20.33 | 65.03 | 13.96 | 88.32 | 167.36 |
|  | significance | *** | ** | *** | ** | *** | *** |

**Table 4.** Phylogenetic signal of variation in muscle volume proportions. Pagel’s *λ* and Blomberg’s K were computed across the full bombycoid tree, pruned to the 20 study species, and within each family on the corresponding subtree. The full-tree signal reflects the deep family-level split, while within-family analyses reveal distinct patterns of trait structuring within each clade. *λ* values close to 1 and K values *>* 1 indicate that variation of the given trait is strongly phylogenetically structured. *λ* p-values from likelihood-ratio tests; K p-values from 1000 randomizations. Significance: *** p < 0.001, ** p < 0.01, * p < 0.05, n.s. p ≥ 0.05.

|  | *Bombycoidea* (n=20) |  | *Saturniidae* (n=10) |  | *Sphingidae* (n=10) |  |
| --- | --- | --- | --- | --- | --- | --- |
| **muscle group** | $\lambda$ | K | $\lambda$ | K | $\lambda$ | K |
| indirect | 0.925*** | 1.19** | 0.846* | 1.18* | 0.547 | 0.80 |
| DLM | 0.800*** | 0.97** | 0.421 | 0.69 | 0.517 | 0.70 |
| DVM | 0.432 | 0.54 | ~0 | 0.61 | ~0 | 0.48 |
| DO | 0.993*** | 3.00** | 1.143*** | 2.73** | 0.149 | 0.47 |
| basalar | ~0 | 0.43 | 0.810 | 0.92* | ~0 | 0.34 |
| subalar | 1.000*** | 3.06** | 1.044* | 1.35** | 0.772 | 0.98* |
| third ax. | 0.993*** | 1.93** | ~0 | 0.56 | 0.617 | 0.83 |

#### **2.2.5** Morphometric analysis

When conducting family comparisons of morphometric parameters, we normalized length and area to the total muscle volume of each animal to remove the effects of body size differences. We divided length and area by the 1/3 and 2/3 exponents of total muscle volume, respectively, to obtain nondimensional normalized values for each.

To assess family-level differences in muscle geometry while accounting for covariance between normalized length and area, we performed two-sample Hotelling’s T^2^ tests for each muscle [Table 3]. Hotelling’s T^2^ tests were performed in R using the hotelling.test() function from the Hotelling package (v1.0-8) (Curran & Hersh, 2012). For each muscle, normalized length and normalized cross-sectional area were treated as a bivariate response variable, and differences between *Sphingidae* and *Saturniidae* were evaluated using Hotelling’s T^2^ statistic. P-values were adjusted for multiple comparisons across the six muscles using Holm correction. For visualization, family centroids were plotted in normalized length-area space and surrounded by 95% confidence ellipses [Fig. 5].

**Figure 5.**
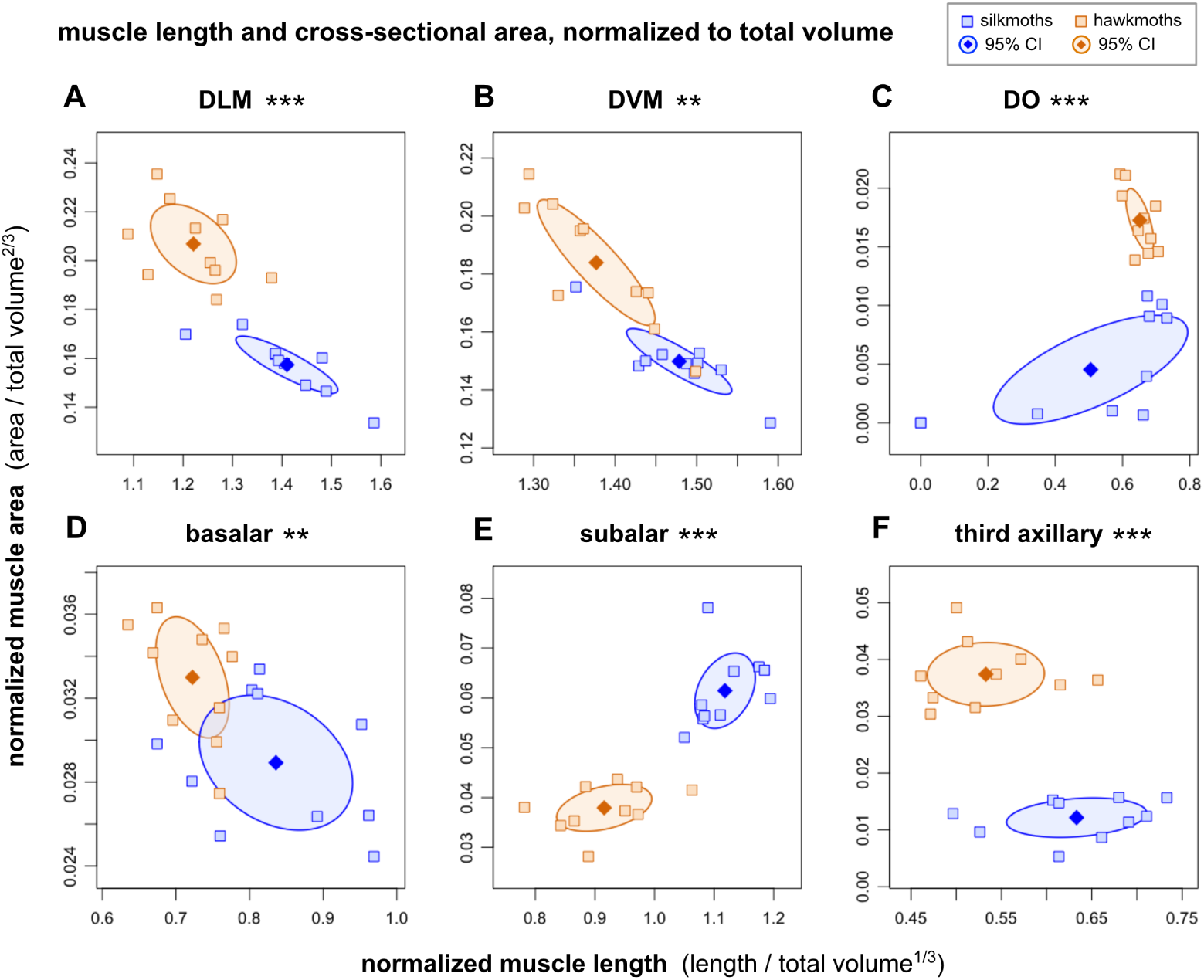
Silkmoths and hawkmoths differ in the geometry of individual flight muscles. Length and area measurements for each muscle are normalized to the 1/3 and 2/3 exponents respectively of the total muscle volume of each specimen. Points represent species, diamonds indicate family centroids, and shaded ellipses represent 95% Hotelling confidence regions around centroid estimates. Asterisks indicate significance of Hotelling’s two-sample T² tests after Holm correction for multiple comparisons (*** p *<* 0.001, ** p *<* 0.01, * p *<* 0.05). **A-F.** Individual muscles show larger normalized lengths in silkmoths and larger normalized areas in hawkmoths, with the exception of the subalar, which is both longer and wider in silkmoths, and the DO, which is both longer and wider in hawkmoths. The third axillary muscle exhibits the largest multivariate divergence in geometry between the families.

**Figure 6.**
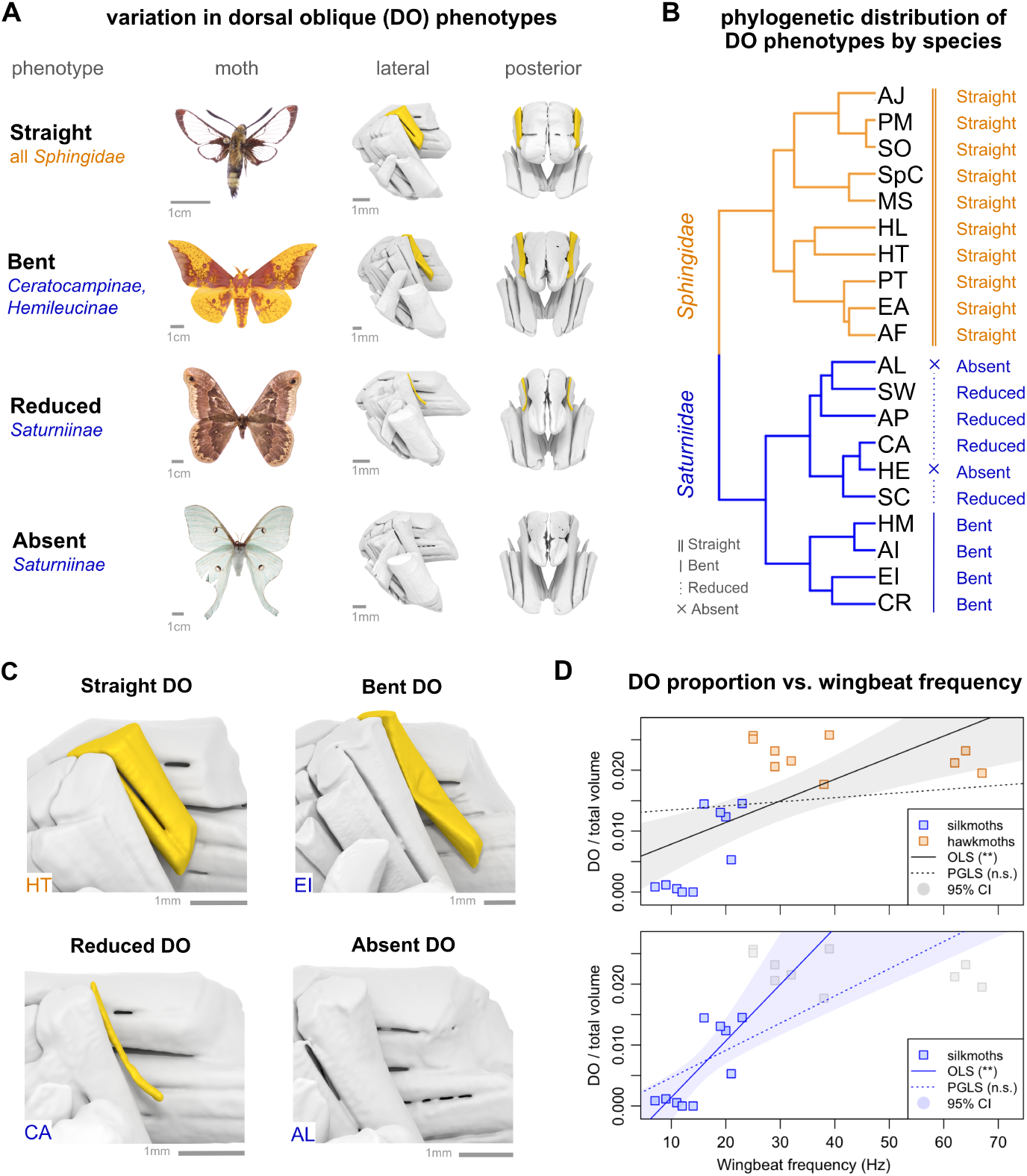
The dorsal oblique (DO) muscle is widely conserved in hawkmoths but morphologically variable in silkmoths. **A.** The DO presents in four phenotypes: Straight, Bent, Reduced, and Absent. These phenotypes are visually discernible and occupy distinct relative volume ranges (body scale bars = 1cm, muscle scale bars = 1mm). **B.** In all ten hawkmoths we observed a Straight DO, while silkmoths had Bent, Reduced, or Absent DO phenotypes. The two silkmoths with an Absent DO are the only specimens observed to have bilateral absence of any flight muscle. The phenotypes of the DO distribute along phylogenetic lines. **C.** Close-up images of the Straight, Bent, Reduced, and Absent phenotypes of the DO (scale bars = 1mm). **D.** DO proportion increases with wingbeat frequency across all sampled bombycoids (top panel) and in silkmoths only (bottom panel). However, these relationships do not remain significant after phylogenetic correction, reflecting the strong phylogenetic structuring of DO proportional volume [Tables 1 & 4]. Among hawkmoths, DO proportion is not significantly correlated with wingbeat frequency, consistent with the relatively uniform DO investment observed across *Sphingidae*.

## **3** Results

### **3.1** Body mass predicts muscle volume

To account for differences in body size across species, we first tested whether total flight muscle volume scales predictably with body mass. Across the full sample, log body mass correlates closely with log total flight muscle volume [Fig. 2C, Table 1]. The strong positive relationship (R^2^ = 0.787, p < 0.001) validates the use of total muscle volume as a normalization for comparing muscle investment across species of different sizes.

The relationship between body mass and total flight muscle volume has a logarithmic scaling exponent of 0.79 (OLS slope = 0.790, PGLS slope = 0.788). An F-test shows that this scaling exponent is significantly hypoallometric compared to the isometric expectation of 1.0 (OLS *F*_1,18_ = 5.04, p = 0.038; PGLS *F*_1,18_ = 5.18, p = 0.035). Thus, larger bombycoid moths possess proportionally less total flight muscle volume than expected under isometric scaling. Examined individually, the DLM and DVM also showed significant hypoallometric scaling; most other muscles also exhibited scaling exponents below 1, but did not deviate significantly from isometric expectations [Supp. Fig. S1, Table S6]. Significant hypoallometry in cross-

### **3.2** Faster-flapping bombycoids have larger proportions of indirect muscle volume

Because indirect and direct flight muscles have distinct functional roles, we tested whether proportional investment in indirect musculature differs between silkmoths and hawkmoths. Hawkmoths allocate a greater proportion of total flight muscle volume to indirect muscles (84.3%, 95% CI: 82.7–85.9%) than silkmoths (79.9%, 95% CI: 77.9–81.8%) [Fig. 3A, Table 2]. This difference is significant when species are treated as independent observations (Welch’s t-test, p < 0.001), but is not significant after phylogenetic correction (phylogenetic ANOVA, p = 0.214). Consistent with this result, indirect muscle proportion exhibits strong phylogenetic signal across *Bombycoidea* (Pagel’s *λ* = 0.925, Blomberg’s K = 1.19) [Table 4]. Indirect muscle proportion is strongly positively correlated with wingbeat frequency (OLS slope = 0.0016 Hz^−1^, R^2^ = 0.704, p < 0.001) [Fig. 3B, Table 1]. The slowest-flapping silkmoth exhibits the lowest indirect muscle proportion (75%), whereas the fastest-flapping hawkmoth exhibits the highest (88%). This relationship remains significant after phylogenetic correction and retains a similar slope (PGLS slope = 0.0012 Hz^−1^, *λ*^^^ = 0.761, p < 0.001), indicating that larger indirect muscle proportions are associated with higher frequencies across lineages despite substantial phylogenetic structuring of the trait.

An OLS regression on silkmoths only shows a substantially steeper slope (0.0043 Hz^−1^) than on hawkmoths only (0.0011 Hz^−1^). ANCOVA reveals a significant wingbeat frequency × clade interaction, confirming that the slope of the relationship between indirect muscle proportion and wingbeat frequency differs between silkmoths and hawkmoths (ANCOVA: OLS p < 0.01, PGLS p < 0.01) [Fig. 3C-D, Table 1].

PGLS correction of the hawkmoth-only regression yields an identical slope and a maximum-likelihood estimate of *λ*^^^ = 0.000 [Table 1], indicating that among hawkmoths, the relationship between indirect muscle proportion and wingbeat frequency is largely independent of phylogenetic relatedness. Within silkmoths, the PGLS slope is slightly reduced relative to the OLS estimate (0.0038 Hz*^−^*^1^) and is accompanied by a moderate estimate of residual phylogenetic covariance (*λ*^^^ = 0.513), but remains significant and substantially steeper than the corresponding hawkmoth relationship [Table 1]. This reduction indicates that part of the indirect muscle-frequency association reflects shared evolutionary history within *Saturniidae*, which also shows significant phylogenetic signal for indirect proportion (*λ* = 0.846, K = 1.18) [Table 4].

Because the dorsal oblique muscle is not known to contribute substantially to flight power generation (Ortega et al., 2019), we repeated these analyses considering only the indirect power muscles (DLM + DVM). The strong positive relationship between indirect muscle allocation and wingbeat frequency was retained, as was significance of the ANCOVA interaction term, indicating that the overall association between frequency and indirect muscle investment is not substantially driven by variation in the dorsal oblique muscle [Supp. Fig. S3]. We thus analyze the three indirect muscle pairs as a group, with their combined proportion exactly inverse to the combined proportion of the direct muscles.

### **3.3** Specific flight muscles diverge in size and morphology between silkmoths and hawkmoths

The observed variation in indirect-direct flight muscle investment raises the question of how individual muscles contribute to this broader pattern. Examination of individual muscles reveals family differences in proportional volume [Fig. 4, Table 2] with corresponding differences in muscle geometry [Fig. 5]. Multivariate comparisons of normalized length and area identify significant family-level differences in all six muscles (Hotelling’s T² = 14.0–167.4; Holm-corrected p < 0.01 in all cases) [Table 3], indicating that silkmoths and hawkmoths occupy distinct regions of muscle morphometric space.

The subalar exhibits one of the strongest family-level differences. Silkmoths devote nearly twice the proportion of total muscle volume to the subalar compared to hawkmoths [Fig. 4B]. The observed proportional divergence remains statistically significant after considering phylogeny [Table 2], despite subalar proportion showing exceptionally strong phylogenetic signal across *Bombycoidea* (Pagel’s *λ* = 1.00, K = 3.06) [Table 4]. Underpinning their enhanced volume, silkmoth subalars are greater in both normalized length and normalized cross-sectional area than hawkmoth subalars [Fig. 5E]. Morphometric analysis of the subalar shows substantial multivariate divergence between families (Hotelling’s T² = 88.3, adjusted p < 0.001) [Table 3], indicating that the enlarged silkmoth subalar reflects coordinated increases in both length and area.

An opposite pattern was observed for the third axillary. Hawkmoths allocate more than twice the proportion of total muscle volume to the third axillary as silkmoths [Fig. 4C]. The volume divergence likewise remains statistically significant after phylogenetic correction [Table 2] despite exhibiting strong phylogenetic signal across *Bombycoidea* (Pagel’s *λ* = 0.993, K = 1.93) [Table 4]. This volume disparity is driven primarily by larger normalized cross-sectional area in hawkmoths, as silkmoths possess longer third axillary muscles [Fig. 5F]. The third axillary exhibits the strongest divergence in geometry among all muscles examined (Hotelling’s T² = 167.4, adjusted p < 0.001), with substantial family-level differences in both normalized length and normalized cross-sectional area.

Basalar muscles do not differ significantly in volume proportion [Fig. 4F, Table 2], but nonetheless occupy distinct regions of morphospace. Hawkmoths exhibit greater normalized cross-sectional area for the basalar, whereas silkmoths exhibit greater normalized length [Fig. 5D]. Although multivariate differences remain significant (Hotelling’s T² = 14.0, adjusted p = 0.008), the basalar shows the weakest family-level geometric divergence among the six muscles examined [Table 3]. The basalar also shows weak phylogenetic signal across *Bombycoidea* and within each family [Table 4].

The indirect power muscles display more subtle divergence patterns. Hawkmoths exhibit larger DLM volume proportions, whereas silkmoths exhibit larger DVM volume proportions [Fig. 4D-E, Table 2]. However, neither difference remains significant under phylogenetic ANOVA, indicating that family-level divergences cannot be distinguished from phylogenetic covariance among species. Despite opposite volumetric trends, both the DLM and DVM follow a similar morphometric pattern, with hawkmoths exhibiting greater normalized cross-sectional area and silkmoths exhibiting greater normalized length [Fig. 5A-B]. Muscle geometry is more differentiated between the families for the DLM (T² = 61.5, adjusted p < 0.001) than the DVM (T² = 20.3, adjusted p = 0.003) [Table 3], consistent with the stronger t-test proportional divergence observed in the DLM.

### **3.4** Dorsal oblique muscles are large and uniform in hawkmoths, but smaller and variable among silkmoths

The dorsal oblique (DO) muscle is conserved in morphology across hawkmoths, but is smaller and sometimes absent in silkmoths [Fig. 4A and 4G]. The average proportion of the DO is over three times larger in hawkmoths (mean = 2.23% of total muscle volume) compared to silkmoths (mean = 0.62%) [Table 2]. The volume disparity appears predominantly driven by hawkmoths having larger normalized DO areas; the DO is also the only muscle in our sample which is not longer in silkmoths [Table 3]. Additionally, DO volume proportions are uniform across hawkmoths (95% CI: 2.04–2.43%; IQR 2.07%-2.46%) but highly labile among silkmoths (95% CI: 0.15–1.09%; IQR 0.06%-1.29%). Underlying this pattern, normalized DO lengths and areas are both tightly clustered in hawkmoths, whereas both metrics include a subset of notably smaller values in silkmoths [Fig. 5C].

We characterized four DO phenotypes which are bilaterally symmetric and distributed across the phylogeny in a structured manner [Fig. 6A-C]. All ten hawkmoths possess a prominent DO occupying 1.5-3.0% of total muscle volume [Fig. 4G]. The consistent DO morphology present in all ten hawkmoth species is termed the Straight DO phenotype due to its muscle fibers appearing uniformly parallel [Fig. 5C], with structural similarity to the highly regular, parallel fiber geometry of the adjacent DVM. The DO appears conserved across *Sphingidae*, suggesting functional importance in hawkmoths.

Silkmoths display three alternative DO phenotypes with distinct volume ranges, all smaller than hawkmoth DOs [Fig. 5A-B]. Four species in the subfamilies *Ceratocampinae* and *Hemileucinae* possess a Bent DO (1.0-1.5% of muscle volume), referring to an indent on the lateral aspect [Fig. 5C]. Four species in the *Saturniinae* subfamily have a Reduced DO (<1%, usually <0.5%) which appears to have far fewer fibers than Straight or Bent DOs. The other two *Saturniinae* silkmoths completely lack the DO. These are the only instances in our study of a flight muscle being absent. These specimens (AL and HE) have a complete lack of muscle tissue in the mesothoracic region where the DO is usually found, while the other five flight muscles are normally developed [Fig. 2A].

The pronounced divergence in DO volume proportion between families remains significant after phylogenetic correction (phylogenetic ANOVA, *p* = 0.046) [Table 2], despite exceptionally strong phylogenetic signal in the trait (Pagel’s *λ* = 0.993, K = 3.00) [Table 4]. Across the full sample, DO proportional volume increases with wingbeat frequency under OLS regression (slope = .0003 Hz^−1^, R^2^ = 0.44, p = 0.001) [Fig. 6D, Table 1]. A similar, steeper positive relationship is observed within *Saturniidae* alone (OLS: .0009 Hz^−1^, R^2^ = 0.60, p = 0.009). However, neither relationship remains significant after phylogenetic correction (pooled PGLS: p = 0.356; *Saturniidae* PGLS: p = 0.083). This loss of significance is underpinned by high phylogenetic covariance of DO proportional volume across *Bombycoidea* (*λ*^^^ = 0.99) and within *Saturniidae* (*λ*^^^ = 1.00) [Table 1].

## **4** Discussion

Corroborating our hypotheses, three primary findings emerge. (1) Indirect muscle proportion is strongly associated with wingbeat frequency, with faster-flapping hawkmoths allocating more volume to indirect musculature than slower-flapping silkmoths. (2) Silkmoths and hawkmoths exhibit family-specific differences in the volume and geometry of individual muscles. Finally, (3) the dorsal oblique muscle is prominent among hawkmoths but diminished to varying degrees among silkmoths, including two instances of absence.

### **4.1** Indirect muscle allocation increases with wingbeat frequency

Indirect muscle allocation is positively correlated with wingbeat frequency in bombycoid moths [Fig. 3], with linear relationships remaining significant after accounting for phylogenetic relatedness [Table 1]. However, the family comparison of indirect proportion loses significance when considering phylogeny [Table 2], so our first hypothesis is partially supported. This suggests that indirect muscle proportion is more closely associated with frequency than with family identity itself. The DLM and DVM are the primary power-producing flight muscles (Pringle, 1949; Tu & Daniel, 2004), and together constitute the vast majority of indirect flight muscle volume, with the dorsal oblique (DO) contributing only a small volume fraction [Fig 4A, Supp. Fig. S3]. Accordingly, larger proportions of indirect muscle suggest more muscle capacity devoted to generating flight power. Mechanical flight power has both inertial and aerodynamic components. Inertial power — the power required to accelerate and decelerate the wings — scales with the cube of frequency (Dickinson et al., 1998; Dudley, 2000); hawkmoths therefore tend to experience greater inertial power demands than silkmoths ( 30-45 W/kg versus 10-15 W/kg) (Aiello, Sikandar, et al., 2021). In contrast, both families converge on similar requirements for aerodynamic power (Aiello, Sikandar, et al., 2021). However, inertial power scaling does not fully explain our data, as steady-state aerodynamic power calculations may not capture peak values of transient aerodynamic demands (Dudley, 2000). Furthermore, thoracic resonance can allow animals to meet high inertial power demands without increasing overall power production (Ellington, 1985).

Resonant mechanics provide crucial context for understanding the observed correlation between wingbeat frequency and indirect muscle proportions. Theoretically, the inertial power costs of high-frequency flight can be canceled out through elastic energy exchange if an insect flaps at its thoracic resonant frequency (Dudley, 2000; Ellington, 1985; Gau et al., 2019). However, hawkmoths forfeit some of this possible energy effiiciency in favor of further enhancing maneuverability. Hawkmoths operate above their thoracic resonant frequency, a phenomenon known as supra-resonance (Gau et al., 2022). This lowers aerodynamic effiiciency compared to flapping at resonance, but makes it easier to perform frequency modulation, a key mechanism hawkmoths utilize for agile maneuvers and stable hovering (Gau et al., 2021). Silkmoths also exhibit supra-resonant flight, but are generally much closer to resonance than hawkmoths (Wold, Aiello, et al., 2024). Because hawkmoths sacrifice effiiciency for maneuverability moreso than silkmoths, they probably rely on larger indirect muscles to meet the compounded demands of high-frequency, supra-resonant flapping. Thus, the positive relationship between indirect muscle allocation and wingbeat frequency likely reflects systematic changes in resonant mechanics across the bombycoid frequency range.

Silkmoths flap slower and closer to their thoracic resonant frequency (Wold, Aiello, et al., 2024), which lowers their inertial costs and presumably reduces demand on the indirect flight power muscles (Dudley, 2000; Ellington, 1985). Likely reflecting their limited energy budget, silkmoths exhibit adaptations such as larger stroke amplitudes, lower wing loading, and passive body oscillations which reduce power demands while contributing to unpredictable flight paths (Aiello, Sikandar, et al., 2021; Aiello, Tan, et al., 2021; Sikandar et al., 2025). Slower wingstrokes may also reduce demands on the indirect flight muscles by providing more time for force development within each stroke cycle (Kammer & Rheuben, 1981; Stevenson & Josephson, 1990). This may allow slower-flapping bombycoids to meet flight power needs with less indirect muscle volume, while deriving greater supplemental power production from direct muscles. Within silkmoths, the association between frequency and indirect muscle proportion is partially influenced by phylogenetic covariance [Fig. 3C, Table 1], likely reflecting frequency range differentiation among silkmoth subfamilies (Janzen, 1984; Sikandar et al., 2025). Both the lowest wingbeat frequencies and the lowest indirect muscle proportions occur in the *Saturniinae* subfamily, whereas the higher-frequency *Hemileucinae* and *Ceratocampinae* have indirect muscle proportions closer to hawkmoths [Figs. 2A & 3C]. In the other direction, the three highest-frequency hawkmoths have the highest indirect muscle proportions, and equivalently the lowest direct muscle proportions [Fig. 3D]. While this follows the expected effects of inertial power scaling and supra-resonance, our data also signal that bombycoids may encounter certain limits at the high frequency extreme. Combined direct muscle proportions do not drop below ~10% in our sample [Table 2, Supp. Table S3], suggesting that the fastest hawkmoths’ direct muscles may approach a minimum volume required to perform essential functions of wing control. Minimum size requirements for direct muscles might therefore contribute to the shallower relationship between frequency and indirect muscle proportion within *Sphingidae* compared to *Saturniidae* [Fig. 3C-D]. At the same time, the DLM is known to supplement steering functionality through asymmetric timing modulation (Sponberg & Daniel, 2012; Sponberg et al., 2015; Wood et al., 2024). This ability likely helps faster moths maintain robust wing control despite heavily prioritizing indirect muscle investment.

Hawkmoths’ higher-frequency flight is important for their ability to hover-feed (Aiello, Sikandar, et al., 2021; Wold, Aiello, et al., 2024), raising the question of whether larger indirect muscles are an adaptation specifically for performing feeding behavior. Our study sample included three non-feeding hawkmoths of the subfamily *Smerinthinae* [Fig. 2A], which have life history and feeding ecology closer to silkmoths, but wingbeat frequency, inertial power demands, and flight kinematics typical of hawkmoths (Aiello, Sikandar, et al., 2021; Aiello et al., 2026; Kawahara, 2007; Tuskes et al., 1996). In *S. ophthalmica*, *P. myops*, and *A. juglandis*, we observe indirect muscle proportions similar to other hawkmoths [Fig. 3D, Supp. Table S3]. Given recent findings that certain head muscles are smaller in *Smerinthinae* than in feeding hawkmoths (Sakagami et al., 2026), *Smerinthinae* appear to maintain large indirect muscles despite lacking other targeted muscular adaptations for feeding. This result further supports that indirect muscle allocation is more closely associated with frequency and flight strategies than with feeding ecology alone. Nevertheless, questions remain about whether the limited energy budget of *Smerinthinae* affects their capacity to perform high-frequency flight with suboptimal energetic effiiciency, suggesting they may have greater reliance on power-reducing wing shapes or elastic energy exchange mechanisms (Aiello, Tan, et al., 2021; Alexander & Bennet-Clark, 1977; Gau et al., 2019).

### **4.2** Families diverge in size and geometry of flight muscles

Examining the volume allocation of individual muscles reveals that the strongest family-level divergences are concentrated in a subset of flight muscles rather than being distributed uniformly throughout the flight apparatus. While t-tests show proportional divergences in five of six muscles [Fig. 4], only the subalar, third axillary, and dorsal oblique remain significantly different after phylogenetic correction [Table 2]. In both size and geometry, these muscles show the strongest and most evolutionarily persistent disparities between silkmoths and hawkmoths, constituting signature features of each family’s musculature. Nevertheless, subtle family-dependent morphological differences appear pervasive across the flight musculature, with all six muscles varying in geometry between the families [Fig. 5]. Multivariate analyses of normalized muscle geometry reveal consistent differences in length versus area scaling, strengthening our overarching conclusion that silkmoths and hawkmoths evolved distinct muscular adaptations for their respective flight strategies.

Subalar muscle investment is a particularly pronounced difference between the families. Silkmoths exhibit substantially larger subalar volume proportions than hawkmoths [Fig. 4B, Table 2], a disparity that arises from greater investment into both length and area [Fig. 5E, Table 3]. The subalar likely controls wing supination; it depresses the trailing edge of the wing during the downstroke, increasing angle of attack and contributing to downstroke force production, and it assists asymmetric wing control during turning (Kammer, 1970, 1971; Ortega et al., 2019; Pfau, 2025). Larger normalized area values suggest that silkmoths have greater relative force capacity in the subalar compared to hawkmoths [Fig. 5]. This relative enhancement may be because their greater stroke amplitude and more inclined stroke planes (Aiello, Sikandar, et al., 2021) require sustained supination of larger wings throughout each wingstroke. The species with the largest subalar investments are in the *Saturniinae* subfamily [Table 4, Supp. Table S3]. Larger subalars may therefore help actuate the behavioral strategies associated with low frequencies, as *Saturniinae* contains the slowest-flapping bombycoids which perform the most pronounced pitching-and-bobbing body oscillations (Sikandar et al., 2025).

Notably, the subalar is the only muscle for which silkmoths have higher normalized cross-sectional area than hawkmoths; there is otherwise a broad trend of hawkmoths having wider muscles while silkmoths have longer muscles. Silkmoths having longer muscles may seem counterintuitive, implying that despite flying at lower frequencies, they evolved muscles with greater potential contraction velocities. However, silkmoths also experience muscle strains roughly twice as large as those of hawkmoths during flight (Wold, Aiello, et al., 2024). Longer fibers permit larger overall muscle excursions while distributing strain across more sarcomeres in series, helping maintain individual sarcomeres within favorable operating ranges (Gordon et al., 1966). Combined with lower-frequency flight, longer muscles may allow silkmoths to operate at a smaller fraction of their maximum shortening velocity, helping maintain favorable force production throughout the wingstroke (Hill, 1938). Thus, silkmoths’ enlargement of the subalar across both area and length suggests prioritization of force for wing supination alongside long fibers to accommodate high muscle strains.

The pattern of hawkmoths having wider muscles while silkmoths have longer muscles is particularly pronounced in the third axillary. Hawkmoths possess substantially larger third axillary volume proportions, driven by greater normalized areas [Figs. 4C & 5F]. The magnitude of the geometric difference exceeds that observed for any other flight muscle, suggesting that the third axillary may have experienced especially divergent functional demands (T^2^ = 167.36, p < 0.001). The third axillary muscles assist wing folding, wing retraction, stroke-plane adjustment, and asymmetric control during turning, playing a key role in pitch and yaw regulation (Ando & Kanzaki, 2004; Kammer, 1971; Pfau, 2025; Rheuben & Kammer, 1987; Wang et al., 2008). Hovering hawkmoths maintain a stable pitch angle across wingstrokes (Aiello, Sikandar, et al., 2021; Hedrick & Daniel, 2006), requiring precise wing and body control which is likely assisted by a robust third axillary muscle. While the role of the third axillary in wing folding is highly consistent across many insects (Kammer, 1971; Pringle, 1968), this result draws attention to how its other known functions may vary in utility between flight strategies.

Family-level differences in basalar size and shape are much weaker than those observed in the subalar or third axillary. Although hawkmoths possess greater normalized basalar area while silkmoths possess greater length [Fig. 5D], basalar volume proportions do not differ significantly between the families [Fig. 4F, Table 2]. The basalars play a primary role in wing pronation, are important for actuating turning, and contribute to wing protraction and stroke-plane adjustment alongside their contribution to force production (Kammer, 1967, 1971; Pfau, 2025). The relatively subtle morphometric differences we observe, alongside no significant disparity in proportion, suggest that the functional demands placed on the basalar are likely more similar between silkmoths and hawkmoths compared to the other two steering muscles. Nevertheless, the larger basalar cross-sectional area in hawkmoths [Fig. 5D] suggests greater investment into pronation force and direct downstroke force.

Examining the indirect power muscles individually reveals a more nuanced pattern than the overall divergence in indirect muscle allocation. Under t-tests, hawkmoths exhibit larger DLM volume proportions and silkmoths exhibit larger DVM volume proportions [Fig. 4D-E], but neither difference remains significant after phylogenetic correction [Table 2]. The family similarity of phylogenetically-corrected DLM and DVM proportions suggests that the indirect power muscle divergence is weaker and less evolutionarily conserved than the pronounced divergences observed in the subalar and third axillary. However, both the DLM and DVM follow the broader geometric trend of hawkmoths having greater normalized cross-sectional areas and silkmoths having greater lengths [Fig. 5A-B]. Accordingly, family divergences are more apparent among the indirect power muscles when examining muscle geometry rather than size alone. While the greater normalized cross-sectional areas of hawkmoths are consistent with increased force-generating capacity, the longer indirect muscles of silkmoths may permit greater shortening excursions while maintaining favorable force-velocity operating conditions.

### **4.3** Dorsal oblique morphology covaries with flight strategy

We observe that the mesothoracic dorsal oblique muscle (DO; elsewhere referred to as II dl2 or the laterophragma muscle (Eaton, 1988; Pfau, 2025)) is profoundly divergent in morphology between the two bombycoid families. Hawkmoths have large DOs with highly uniform geometry [Figs. 4G & 5C], whereas silkmoths have smaller DOs that are unusually labile in both size and morphology [Fig. 6]. When animal lineages evolve increasingly specialized ecological and behavioral strategies, traits that no longer benefit functional performance can accumulate phenotypic variation and eventually become reduced or lost (Higham et al., 2015; Wilkens, 2021). This is a general evolutionary principle which extends specifically to insect flight musculature (Lu et al., 2020; Marden, 2000). The smaller DO phenotypes present among silkmoths, including two instances of complete absence, appear consistent with lessened biomechanical demand permitting de-investment in this muscle. In contrast, the large DOs present across all sampled *Sphingidae* suggest sustained functional importance for the DO throughout hawkmoth diversification.

Our findings expand upon the observation of Pringle (1968) that “all species known to possess [a DO] can hover.” We find that most silkmoths possess some smaller form of the muscle despite not hovering; however, the reduction of the silkmoth DO relative to hawkmoths is consistent with silkmoths facing less selective pressure to allocate volume for the DO. We also find that the DO appears important for hawkmoth flight even in species which do not hover-feed, mirroring the broader pattern observed for indirect flight muscle investment. The *Smerinthinae* in our sample [Fig. 2A] — which are non-feeding as adults, but retain hawkmoth-typical wingbeat frequencies and flight kinematics (Aiello, Sikandar, et al., 2021; Kitching & Cadiou, 2000) — each possess robust Straight DOs [Fig. 6B]. A large DO therefore appears to be an integrated component of the hawkmoth flight apparatus, associated with a broader suite of adaptations for high-frequency, maneuverable flight. However, *Sphingidae* alone show no significant association between wingbeat frequency and DO pro- portion. Furthermore, although DO investment increases with frequency across bombycoids overall, this relationship loses significance after phylogenetic correction [Fig. 6D, Table 1]. These findings suggest that DO investment varies between lineages specialized for different flight strategies rather than scaling continuously with frequency.

This interpretation is supported by the distribution of DO phenotypes among the silkmoth phylogeny because the two silkmoth subclades in our sample show different degrees of DO investment aligning with known contrasts in flight frequency and behavior. Reduced and Absent DO phenotypes occur exclusively within the *Saturniinae*, which contain the lowest-frequency silkmoths in our sample and represent some of the largest known moths [Figs. 2A, 6A-B]. Notably, *S. walterorum*, the only sampled *Saturniinae* exceeding ~15 Hz, possesses the largest DO within that subfamily [Supp. Table S3]. In contrast, the *Hemileucinae* and *Ceratocampinae* possess larger Bent DOs [Fig. 6] and tend to have more hawkmoth-like wingbeat frequencies and flight kinematics than *Saturniinae* (Janzen, 1984; Sikandar et al., 2025; Wold, Aiello, et al., 2024). OLS regression identifies a positive association between DO investment and wingbeat frequency among silkmoths alone, but as in the pooled bombycoid analysis, significance diminishes after phylogenetic correction, with *λ*^^^ approaching 1 [Fig. 6D, Table 1]. Thus, instead of scaling independently with wingbeat frequency, DO investment appears closely associated with lineage identity, with larger DOs occurring in clades that generally occupy moderate-to-high portions of the bombycoid frequency spectrum. These results also strengthen evidence that the *Saturniinae* and *Hemileucinae*-*Ceratocampinae* silkmoth lineages possess distinct integrated biomechanical strategies that stem from their early evolutionary divergence.

One function proposed for the dorsal oblique is physical modulation of the elastic exoskeleton (Ortega et al., 2019), which could explain the muscle’s conservation in hawkmoths and higher-frequency silkmoths. The DO originates dorsally on the scutum (Eaton, 1988), a critical site of exoskeletal deformation by the DLM and DVM (Ando & Kanzaki, 2016; Gau et al., 2019). This attachment position is consistent with the DO influencing how strains generated by the DLM and DVM propagate through the thoracic exoskeleton to the wing hinge, providing a potential mechanism for indirect modulation of wing motion and flight control. In *Diptera*, the pleurosternal muscle is thought to tune resonant mechanics by altering exoskeletal stiffness (Dickinson & Tu, 1997; Nachtigall & Wilson, 1967), and the lepidopteran DO has been speculated to play a similar role (Ortega et al., 2019). Since resonant mechanics and behavioral strategy both vary systematically with wingbeat frequency among *Bombycoidea* (Aiello, Sikandar, et al., 2021; Wold, Aiello, et al., 2024; Wold, Liu, et al., 2024), frequency would likely influence the usefulness of this hypothetical mechanism. In higher-frequency hawkmoths which flap further from resonance and have sharper resonant curves (Wold, Aiello, et al., 2024), flight performance may be more sensitive to subtle alterations of thoracic stiffness and strain propagation. A muscle which modulates these properties may be more valuable to these species than to slower-flapping silkmoths, which are already more aerodynamically effiicient and whose behavioral strategies do not emphasize precise control of body position (Aiello, Sikandar, et al., 2021; Wold, Aiello, et al., 2024).

The DO is also sometimes categorized as an wing elevator muscle, given its location adjacent and parallel to the DVMs (Dudley, 2000; Kammer, 1967; Pfau, 2025). The DO is known to fire in phase with the DVM, the expected pattern for a wing elevator, but the studies which record this also report substantial variation in DO activation timing that suggest a more complex role (Kammer, 1970, 1971; Ortega et al., 2019). The DO may contribute to upstroke mechanics beyond the action of the DVMs, potentially extending upstroke range or increasing wing supination during the upstroke (Pfau, 2025; Pringle, 1968). If so, the DO could influence torques generated near the upstroke-downstroke transition, possibly facilitating amplitude modulation and/or pitch stabilization (Gau et al., 2021). Notably, the silkmoths that Sikandar et al. (2025) found to rely most heavily on body pitch oscillations all possess either a Reduced or Absent DO, whereas *A. io* possesses a Bent DO and flies with a comparatively stable pitch angle. While correlative, this pattern could be consistent with the DO actively countering pitch oscillations. Finally, the opposing patterns of investment seen in the DO versus the subalar are intriguing in light of recent work suggesting that the DO and subalar may operate antagonistically on certain associated thoracic structures (Pfau, 2025). Though the DO’s actual function(s) remain unresolved, our data signal that a large DO confers an advantage to hawkmoths and higher-frequency silkmoths which does not seem beneficial to lower-frequency silkmoths, opening compelling opportunities for future functional investigations.

## **5** Conclusion

We demonstrate that silkmoths and hawkmoths possess distinct patterns of flight muscle allocation that correspond closely to their contrasting flight strategies. Wingbeat frequency emerges as a major predictor of how volume is distributed between the power and steering muscles responsible for core flight functionality. Flight muscle evolution is non-uniform, with some muscles exhibiting more pronounced divergences in size and geometry than others. Moreover, these divergences do not all follow the same pattern: indirect muscle allocation is more strongly associated with wingbeat frequency than with lineage identity, whereas several individual muscles exhibit size differences which are closely tied to evolutionary history. Nevertheless, this small suite of muscles appears capable of supporting substantial kinematic diversification with only subtle morphological tuning. This makes the extreme phenotypic divergence of the dorsal oblique muscle particularly notable, motivating further investigation of indirect control mechanisms in the elastic thorax. Among bombycoid moths, flight muscle allocation parallels known frequency-associated variation in wing morphology, thorax mechanics, and behavior. These results further strengthen the utility of *Bombycoidea* as a model clade for studying how integrated locomotor systems evolve, and highlight the value of the broad frequency range represented across silkmoths and hawkmoths (Aiello et al., 2026). More broadly, we present evidence that insect thoracic muscle allocation is evolutionarily sensitive and can adapt alongside external traits to support diversification of locomotor strategies.

## **7** Data Availability

Supplementary Information will be available electronically. Digital 3D renderings of flight muscle morphology from µCT data will be available on MorphoSource.

## **8** Ethics

The authors declare no conflicts of interest.

## **9** Declaration of Generative AI Usage

In preparing this manuscript, the authors employed generative artificial intelligence to support code writing for graphs and statistical analyses, and to refine wording and idea organization. This tool was used with careful oversight, and all analyses, interpretations, and conclusions are fully the work of the authors.

## **10** Author Contributions

Conceptualization: B.A., E.W., J.B., L.W., S.S.; Methodology: B.A., E.W., J.B., L.W., S.S.; Investigation: E.W., J.B.; Software: E.W., J.B., L.W.; Formal Analysis: E.W., J.B.; Resources: B.A., E.W., L.W., S.S.; Writing - original draft preparation: J.B.; Writing - review and editing: B.A., E.W., J.B., L.W., S.S.; Visualization: J.B.; Supervision: E.W., S.S.; Project administration: E.W., S.S.; Funding acquisition: S.S.

## Acknowledgements

The photo in Fig. 1B of *M. sexta* in flight was taken by Rob Felt. All other photos of moths were taken by Brett Aiello.

The authors thank current and former members of the Agile Systems Lab at Georgia Tech for assistance and feedback on this research.

## References

1. Agosta, S. J., & Janzen, D. H. (2005). Body size distributions of large Costa Rican dry forest moths and the underlying relationship between plant and pollinator morphology. Oikos, 108(1), 183–193. 10.1111/j.0030-1299.2005.13504.x

2. Aiello, B. R., Baker, J. N., Barber, J., Joseph, J. J., Kawahara, A. Y., Ketler, G. M., Mowery, S. G., Sikandar, U. B., Sponberg, S., Laurent, R. A. S., Wold, E. S., & Wood, L. (2026). Bombycoid moths: An emerging model clade for studying insect flight. *Integrative And Comparative Biology*, icag143. 10.1093/icb/icag143

3. Aiello, B. R., Sikandar, U. B., Minoguchi, H., Bhinderwala, B., Hamilton, C. A., Kawahara, A. Y., & Sponberg, S. (2021). The evolution of two distinct strategies of moth flight. Journal of The Royal Society Interface, 18(185), 20210632. 10.1098/rsif.2021.0632

4. Aiello, B. R., Tan, M., Bin Sikandar, U., Alvey, A. J., Bhinderwala, B., Kimball, K. C., Barber, J. R., Hamilton, C. A., Kawahara, A. Y., & Sponberg, S. (2021). Adaptive shifts underlie the divergence in wing morphology in bombycoid moths. Proceedings of the Royal Society B: Biological Sciences, 288(1956), 20210677. 10.1098/rspb.2021.0677

5. Alexander, R. M., & Bennet-Clark, H. C. (1977). Storage of elastic strain energy in muscle and other tissues. Nature, 265(5590), 114–117. 10.1038/265114a0

6. Ando, N., & Kanzaki, R. (2004). Changing Motor Patterns of the 3rd Axillary Muscle Activities Associated with Longitudinal Control in Freely Flying Hawkmoths. Zoological Science, 21(2), 123–130. 10.2108/zsj.21.123

7. Ando, N., & Kanzaki, R. (2016). Flexibility and control of thorax deformation during hawkmoth flight. Biology Letters, 12(1), 20150733. 10.1098/rsbl.2015. 0733

8. Camargo, W. R. F. D., Camargo, N. F. D., Corrêa, D. D. C. V., Camargo, A. J. A. D., & Diniz, I. R. (2015). Sexual Dimorphism and Allometric Effects Associated With the Wing Shape of Seven Moth Species of Sphingidae (Lepidoptera: Bombycoidea). Journal of Insect Science, 15(1), 107. 10.1093/jisesa/iev083

9. Chapman, R. F. (1998). The insects: Structure and function (4th ed). Cambridge University Press.

10. Curran, J., & Hersh, T. (2012, October). Hotelling: Hotelling’s t^2^*testandvariants*. 10.32614/CRAN.package.Hotelling

11. Deakin, M. A. B. (2010). Formulae for Insect Wingbeat Frequency. Journal of Insect Science, 10, 96. 10.1673/031.010.9601

12. Dickinson, M. H., Lehmann, F.-O., & Chan, W. P. (1998). The control of mechanical power in insect flight. American Zoologist, 38(4), 718–728. 10.1093/icb/ 38.4.718

13. Dickinson, M. H., & Tu, M. S. (1997). The Function of Dipteran Flight Muscle. Comparative Biochemistry and Physiology Part A: Physiology, 116(3), 223–238. 10.1016/S0300-9629(96)00162-4

14. Dudley, R. (2000). The Biomechanics of Insect Flight: Form, Function, Evolution. Princeton University Press.

15. Eaton, J. L. (1988). Lepidopteran anatomy. Wiley.

16. Ellington, C. P. (1985). Power and effiiciency of insect flight muscle. Journal of Experimental Biology, 115(1), 293–304. 10.1242/jeb.115.1.293

17. Farina, S. C., Kane, E. A., & Hernandez, L. P. (2019). Multifunctional structures and multistructural functions: Integration in the evolution of biomechanical systems. Integrative and Comparative Biology, 59(2), 338–345. 10.1093/icb/ icz095

18. Fedorov, A., Beichel, R., Kalpathy-Cramer, J., Finet, J., Fillion-Robin, J.-C., Pujol, S., Bauer, C., Jennings, D., Fennessy, F., Sonka, M., Buatti, J., Aylward, S., Miller, J. V., Pieper, S., & Kikinis, R. (2012). 3D Slicer as an image computing platform for the Quantitative Imaging Network. Magnetic Resonance Imaging, 30(9), 1323–1341. 10.1016/j.mri.2012.05.001

19. Garland, T., Dickerman, A. W., Janis, C. M., & Jones, J. A. (1993). Phylogenetic analysis of covariance by computer simulation. Systematic Biology, 42(3), 265–292. 10.1093/sysbio/42.3.265

20. Gau, J., Gemilere, R., (Fm Subteam), L.-V., Lynch, J., Gravish, N., & Sponberg, S. (2021). Rapid frequency modulation in a resonant system: Aerial perturbation recovery in hawkmoths. Proceedings of the Royal Society B: Biological Sciences, 288(1951), 20210352. 10.1098/rspb.2021.0352

21. Gau, J., Gravish, N., & Sponberg, S. (2019). Indirect actuation reduces flight power requirements in *Manduca sexta* via elastic energy exchange. Journal of The Royal Society Interface, 16(161), 20190543. 10.1098/rsif.2019.0543

22. Gau, J., Wold, E. S., Lynch, J., Gravish, N., & Sponberg, S. (2022). The hawkmoth wingbeat is not at resonance. Biology Letters, 18(5), 20220063. 10.1098/rsbl. 2022.0063

23. Gordon, A. M., Huxley, A. F., & Julian, F. J. (1966). The variation in isometric tension with sarcomere length in vertebrate muscle fibres. The Journal of Physiology, 184(1), 170–192. 10.1113/jphysiol.1966.sp007909

24. Grafen. (1989). The phylogenetic regression. *Philosophical Transactions of the Royal Society of London. B*, Biological Sciences, 326(1233), 119–157. 10.1098/ rstb.1989.0106

25. Hedrick, T. L., & Daniel, T. L. (2006). Flight control in the hawkmoth manduca sexta: The inverse problem of hovering. Journal of Experimental Biology, 209(16), 3114–3130. 10.1242/jeb.02363

26. Higham, T. E., Birn-Jeffery, A. V., Collins, C. E., Hulsey, C. D., & Russell, A. P. (2015). Adaptive simplification and the evolution of gecko locomotion: Morphological and biomechanical consequences of losing adhesion. Proceedings of the National Academy of Sciences, 112(3), 809–814. 10.1073/pnas.1418979112

27. Hill, A. V. (1938). The heat of shortening and the dynamic constants of muscle. Proceedings of the Royal Society of London. Series B - Biological Sciences, 126(843), 136–195. 10.1098/rspb.1938.0050

28. Humphries, D. A., & Driver, P. M. (1970). Protean defence by prey animals. Oecologia, 5(4), 285–302. 10.1007/BF00815496

29. Janzen, D. H. (1984). Two ways to be a big tropical moth: Santa Rosa saturniids and sphingids. Oxford Surveys in Evolutionary Biology, 1. https://www.researchgate.net/ profile/Daniel-Janzen-2/publication/283604644_Two_ways_to_be_a_tropical_big_moth_Santa_Rosa_saturniids_and_sphingids/links/565c284008aefe619b251e65/ Two-ways-to-be-a-tropical-big-moth-Santa-Rosa-saturniids-and-sphingids.pdf

30. Josephson, R. K. (1999). Dissecting muscle power output. Journal of Experimental Biology, 202(23), 3369–3375. 10.1242/jeb.202.23.3369

31. Kammer, A. E. (1967). Muscle Activity During Flight in Some large Lepidoptera. Journal of Experimental Biology, 47 (2), 277–295. 10.1242/jeb.47.2.277

32. Kammer, A. E. (1970). A comparative study of motor patterns during pre-flight warm-up in hawkmoths. Zeitschrift für vergleichende Physiologie, 70(1), 45–56. 10.1007/BF00299536

33. Kammer, A. E. (1971). The motor output during turning flight in a hawkmoth, *Manduca sexta*. Journal of Insect Physiology, 17 (6), 1073–1086. 10.1016/0022-1910(71)90011-4

34. Kammer, A. E., & Rheuben, M. B. (1981). Neuromuscular mechanisms of insect flight. In C. F. Herreid & C. R. Fourtner (Eds.), Locomotion and energetics in arthropods (pp. 163–194). Springer US. 10.1007/978-1-4684-4064-5_7

35. Kawahara, A. Y. (2007). Molecular phylogenetic analysis of the hawkmoths (lepidoptera: Bombycoidea: Sphingidae) and the evolution of the sphingid proboscis [Copyright - Database copyright ProQuest LLC; ProQuest does not claim copyright in the individual underlying works; Last updated - 2023-03-02]. ProQuest Dissertations and Theses, 159. https://www.proquest.com/dissertations-theses/molecular-phylogenetic-analysis-hawkmoths/docview/304850997/se-2

36. Kawahara, A. Y., & Barber, J. R. (2015). Tempo and mode of antibat ultrasound production and sonar jamming in the diverse hawkmoth radiation. Proceedings of the National Academy of Sciences, 112(20), 6407–6412. 10.1073/pnas. 1416679112

37. Kitching, I. J., & Cadiou, J. M. (2000). Hawkmoths of the world: An annotated and illustrated revisionary checklist (Lepidoptera: Sphingidae). Natural History Museum.

38. Lasso, A. (2025, June). Lassoan/SlicerSegmentEditorExtraEffects. Retrieved July 22, 2025, from https://github.com/lassoan/SlicerSegmentEditorExtraEffects

39. Lieber, R. L., & Ward, S. R. (2011). Skeletal muscle design to meet functional demands. Philosophical Transactions of the Royal Society B: Biological Sciences, 366(1570), 1466–1476. 10.1098/rstb.2010.0316

40. Lindsay, T., Sustar, A., & Dickinson, M. (2017). The function and organization of the motor system controlling flight maneuvers in flies. Current Biology, 27 (3), 345–358. 10.1016/j.cub.2016.12.018

41. Lu, K., Liang, S., Han, M., Wu, C., Song, J., Li, C., Wu, S., He, S., Ren, J., Hu, H., Shen, J., Tong, X., & Dai, F. (2020). Flight muscle and wing mechanical properties are involved in flightlessness of the domestic silkmoth, bombyx mori. Insects, 11(4), 220. 10.3390/insects11040220

42. Marden, J. H. (2000). Variability in the Size, Composition, and Function of Insect Flight Muscles. Annual Review of Physiology, 62(1), 157–178. 10.1146/ annurev.physiol.62.1.157

43. Martin, M. L., Travouillon, K. J., Fleming, P. A., & Warburton, N. M. (2020). Review of the methods used for calculating physiological cross-sectional area (PCSA) for ecological questions. Journal of Morphology, 281(7), 778–789. 10.1002/jmor.21139

44. Nachtigall, W., & Wilson, D. M. (1967). Neuro-Muscular Control of Dipteran Flight. Journal of Experimental Biology, 47 (1), 77–97. 10.1242/jeb.47.1.77

45. O, J., Kwon, H.-J., Kim, S. H., Cho, T.-H., & Yang, H.-M. (2019). Use of micro x-ray computed tomography with phosphotungstic acid preparation to visualize human fibromuscular tissue. Journal of Visualized Experiments: JoVE, (151). https://doi. org/10.3791/59752

46. Orsbon, C. P., Gidmark, N. J., & Ross, C. F. (2018). Dynamic Musculoskeletal Functional Morphology: Integrating diceCT and XROMM. The Anatomical Record, 301(2), 378–406. 10.1002/ar.23714

47. Ortega, J., Angjelichinoski, M., Ravier, R., Ferrari, S., Tarokh, V., & Sponberg, S. (2021, July). Consistent coordination patterns provide near perfect behavior decoding in a comprehensive motor program for insect flight. 10.1101/2021.07. 13.452211

48. Ortega, J., Conn, R., & Sponberg, S. (2019). Precise timing is ubiquitous, consistent, and coordinated across a comprehensive, spike-resolved flight motor program. Proceedings of the National Academy of Sciences, 116(52), 26951–26960. 10.1073/pnas.1907513116

49. Ortega, J., Niebur, T., Wood, L., Conn, R., & Sponberg, S. (2023). An information theoretic method to resolve millisecond-scale spike timing precision in a comprehensive motor program (P. E. Latham, Ed.). PLOS Computational Biology, 19(6), e1011170. 10.1371/journal.pcbi.1011170

50. Pearse, W. D., Davies, T. J., & Wolkovich, E. M. (2023). How to define, use, and interpret pagel’s (lambda) in ecology and evolution. 10.1101/2023.10.10. 561651

51. Pennycuick, C. J., & Rezende, M. A. (1984). The Specific Power Output Of Aerobic Muscle, Related To The Power Density Of Mitochondria. Journal of Experimental Biology, 108(1), 377–392. 10.1242/jeb.108.1.377

52. Pfau, H. K. (2025). Functional Morphology of the Mesothoracic Flight Apparatus of Manduca sexta (Lepidoptera, Sphingidae). 10.13140/RG.2.2.12902.89924

53. Pringle, J. W. S. (1949). The excitation and contraction of the flight muscles of insects. The Journal of Physiology, 108(2), 226–232. 10.1113/jphysiol.1949. sp004326

54. Pringle, J. (1968). Comparative Physiology of the Flight Motor. In Advances in Insect Physiology (pp. 163–227, Vol. 5). Elsevier. 10.1016/S0065-2806(08) 60229-5

55. Revell, L. (2013). R: Compute phylogenetic signal with two methods. Retrieved August 13, 2025, from http://www.phytools.org/static.help/phylosig.html

56. Rheuben, M. B., & Kammer, A. E. (1987). Structure and innervation of the third axillary muscle of *Manduca* relative to its role in turning flight. Journal of Experimental Biology, 131(1), 373–402. 10.1242/jeb.131.1.373

57. Roberts, T. J., Eng, C. M., Sleboda, D. A., Holt, N. C., Brainerd, E. L., Stover, K. K., Marsh,

58. R. L., & Azizi, E. (2019). The multi-scale, three-dimensional nature of skeletal muscle contraction. Physiology, 34(6), 402–408. 10.1152/physiol. 00023.2019

59. Rome, L. C., & Lindstedt, S. L. (1998). The quest for speed: Muscles built for high-frequency contractions. Physiology, 13(6), 261–268. 10.1152/physiologyonline. 1998.13.6.261

60. Sakagami, K., Hamaguchi, K., Terada, K., Takami, Y., & Sugiura, S. (2026). Relationship between proboscis length and dilator muscle volume in japanese hawkmoths (lepidoptera: Sphingidae). Entomological Science, 29(2), e70023. 10.1111/ens.70023

61. Sikandar, U. B., Aiello, B. R., & Sponberg, S. (2025). Body oscillations couple with wing flapping to reduce aerodynamic power in wild silk moth flight. 10.1098/rsif.2025.0061

62. Sponberg, S., & Daniel, T. L. (2012). Abdicating power for control: A precision timing strategy to modulate function of flight power muscles. Proceedings of the Royal Society B: Biological Sciences, 279(1744), 3958–3966. 10.1098/ rspb.2012.1085

63. Sponberg, S., Daniel, T. L., & Fairhall, A. L. (2015). Dual Dimensionality Reduction Reveals Independent Encoding of Motor Features in a Muscle Synergy for Insect Flight Control (O. Sporns, Ed.). PLOS Computational Biology, 11(4), e1004168. 10.1371/journal.pcbi.1004168

64. Springthorpe, D., Fernández, M. J., & Hedrick, T. L. (2012). Neuromuscular control of free-flight yaw turns in the hawkmoth manduca sexta. Journal of Experimental Biology, 215(10), 1766–1774. 10.1242/jeb.067355

65. Stevenson, R. D., & Josephson, R. K. (1990). Effects of operating frequency and temperature on mechanical power output from moth flight muscle. Journal of Experimental Biology, 149(1), 61–78. 10.1242/jeb.149.1.61

66. Swart, P., Wicklein, M., Sykes, D., Ahmed, F., & Krapp, H. G. (2016). A quantitative comparison of micro-ct preparations in dipteran flies. Scientific Reports, 6(1), 39380. 10.1038/srep39380

67. Symonds, M. R. E., & Blomberg, S. P. (2014). A Primer on Phylogenetic Generalised Least Squares. In L. Z. Garamszegi (Ed.), Modern Phylogenetic Comparative Methods and Their Application in Evolutionary Biology: Concepts and Practice (pp. 105–130). Springer. 10.1007/978-3-662-43550-2_5

68. Tu, M. S., & Daniel, T. L. (2004). Submaximal power output from the dorsolongitudinal flight muscles of the hawkmoth *Manduca sexta*. Journal of Experimental Biology, 207 (26), 4651–4662. 10.1242/jeb.01321

69. Tuskes, P. M., Tuttle, J. P., & Collins, M. M. (1996). The Wild Silk Moths of North America: A Natural History of the Saturniidae of the United States and Canada [Google-Books-ID: 3vqpGATXU2oC]. Cornell University Press.

70. Wainwright, P. C., & Price, S. A. (2016). The impact of organismal innovation on functional and ecological diversification. Integrative and Comparative Biology, 56(3), 479–488. 10.1093/icb/icw081

71. Wang, H., Ando, N., & Kanzaki, R. (2008). Active control of free flight manoeuvres in a hawkmoth, *Agrius convolvuli*. Journal of Experimental Biology, 211(3), 423–432. 10.1242/jeb.011791

72. Wilkens, H. (2021). Variability and the primacy of the genotype. Biological Journal of the Linnean Society, 133(4), 931–948. 10.1093/biolinnean/blab065

73. Willmott, A. P., & Ellington, C. P. (1997). The mechanics of flight in the hawkmoth *Manduca sexta* II. Aerodynamic consequences of kinematic and morphological variation. Journal of Experimental Biology, 200(21), 2723–2745. 10.1242/jeb. 200.21.2723

74. Woittiez, R. D., Huijing, P. A., Boom, H. B. K., & Rozendal, R. H. (1984). A three-dimensional muscle model: A quantified relation between form and function of skeletal muscles. Journal of Morphology, 182(1), 95–113. 10.1002/jmor.1051820107

75. Wold, E. S., Aiello, B., Harris, M., Bin Sikandar, U., Lynch, J., Gravish, N., & Sponberg, S. (2024). Moth resonant mechanics are tuned to wingbeat frequency and energetic demands. Proceedings of the Royal Society B: Biological Sciences, 291(2025), 20240317. 10.1098/rspb.2024.0317

76. Wold, E. S., Liu, E., Lynch, J., Gravish, N., & Sponberg, S. (2024). The Weis-Fogh Number Describes Resonant Performance Tradeoffs in Flapping Insects. Integrative And Comparative Biology, 64(2), 632–643. 10.1093/icb/icae039

77. Wood, L. J., Putney, J., & Sponberg, S. (2024). Flight power muscles have a coordinated, causal role in controlling hawkmoth pitch turns. Journal of Experimental Biology, 227 (24), jeb246840. 10.1242/jeb.246840

78. Yang, H., Putney, J., Bin Sikandar, U., Zhu, P., Sponberg, S., & Ferrari, S. (2022). A Relative Spike-Timing Approach to Kernel-Based Decoding Demonstrated for Insect Flight Experiments. 2022 International Joint Conference on Neural Networks (IJCNN), 1–7. 10.1109/IJCNN55064.2022.9892352

79. Yu, W., Zhou, Y., Guo, J., Wyckhuys, K. A. G., Shen, X., Li, X., Ge, S., Liu, D., & Wu, K. (2020). Interspecific and Seasonal Variation in Wingbeat Frequency Among Migratory Lepidoptera in Northern China (Y. Gao, Ed.). Journal of Economic Entomology, 113(5), 2134–2140. 10.1093/jee/toaa134

