## Supplementary Information for "Flight muscle allocation diverges between two moth families with distinct flight strategies"

### SI.1 Protocol for µCT scan segmentation

To segment flight muscles from the µCT scans, we used 3D Slicer (v. 5.6.2) (Fedorov et al., 2012) with the Slicer Extra Effects package (Lasso, 2025). The following manual segmentation protocol allows for clean isolation of muscle geometries while preserving fine structural features.

**Shell** Using the *Surface Cut* tool from Extra Effects on the "fill inside" and "allow overlap" modes, we traced a 3D shell enclosing the entire area of the muscle.

**Threshold** We isolated muscle tissue from surrounding space using the *Threshold* tool at an appropriate intensity for the scan contrast. We set Masking → Editable Area inside the selected segment and then reset the editable area after thresholding.

**Trim** In the *Islands* tool, we applied Keep Largest Island. We then used the *Scissor* tool in the 3D view to manually trim off any remaining extraneous tissue, including edges of adjacent muscles.

**Smooth** In the *Smoothing* tool, we used the following protocol. Exact values slightly varied for particularly large or small scans.

1. Closing (fill holes) at 0.050mm/3x3px
2. Opening (remove extrusions) at 0.050mm/3x3px
3. Threshold with editable area as segment; reset editable area
4. Median at 0.050mm/3x3px
5. Logical operators, to remove overlaps with adjacent muscles
6. Closing (fill holes) at 0.070mm/5x5px
7. Gaussian at s.d. 0.030mm

Following segmentation, we exported 3D .obj files to Blender for rendering.

### SI.2 Hypometric scaling of muscle volume and body mass

We find that bombycoids have static allometry (Stern & Emlen, 1999) between muscle volume and body mass. This pattern extends across *Saturniidae* and *Sphingidae* alike, which both exhibit wide variation in body size, even between closely-related species. According to our result in 3.1, total flight muscle volume scales hypometrically with body mass by a factor of around $M^{0.8}$, which remains after accounting for phylogenetic relationships. In other words, for any bombycoid, a given increase in body mass corresponds to a slightly lesser increase in muscle volume than expected at isometry ($M^1$). Analysis of individual muscles reveals similar hypoallometric scaling in the two largest mesothoracic flight muscles, the dorsolongitudinal muscle (DLM) and dorsoventral muscle (DVM); both scale with slopes significantly below the isometric expectation of 1, including after phylogenetic correction [Fig. S1]. The other four muscles also exhibit regression slopes below 1, but these deviations were not statistically distinguishable from isometry by an F-test [Table S6].

Our observation of hypoallometry is contextualized by previous findings from Bartholomew and Casey (1978) that oxygen consumption scales hypometrically with body mass in both *Sphingidae* and *Saturniidae*. According to this study, bombycoid resting metabolic rate (RMR) scales to body mass by a factor of 0.775, while active metabolic rate (AMR) scales by a factor of 0.818. AMR, which can be orders of magnitude higher than RMR in insects (Beenakkers et al., 1984), is particularly linked with muscle size (Niven & Scharlemann, 2005; Tolfrey et al., 2006; Weibel, 2002) because muscles account for the vast majority of oxygen consumption during flight (Ellington, 1985; Suarez, 2000). Though a causal relationship cannot be inferred, this data aligns closely with our observed scaling of muscle volume and body mass.

Possible explanations for hypometric muscle volume scaling could relate to smaller insects having greater mass-specific power requirements, supported by research in *Sphingidae* (Casey, 1981). On the other hand, larger insects likely face tighter geometric and metabolic constraints on overall thorax size, particularly due to limitations on the diffusive capacity of the tracheal respiratory system (Dudley, 2000). Yet despite over a century of empirical observations that animal metabolic rates follow hypometric scaling (Hulbert, 2014; Kleiber, 1932), current evidence still does not converge on a universal scaling law or a single causative factor for metabolic hypoallometry (Harrison, 2018; Harrison et al., 2022; White et al., 2007). Hypometric scaling of muscle volume may similarly reflect the combined effect of multiple factors.

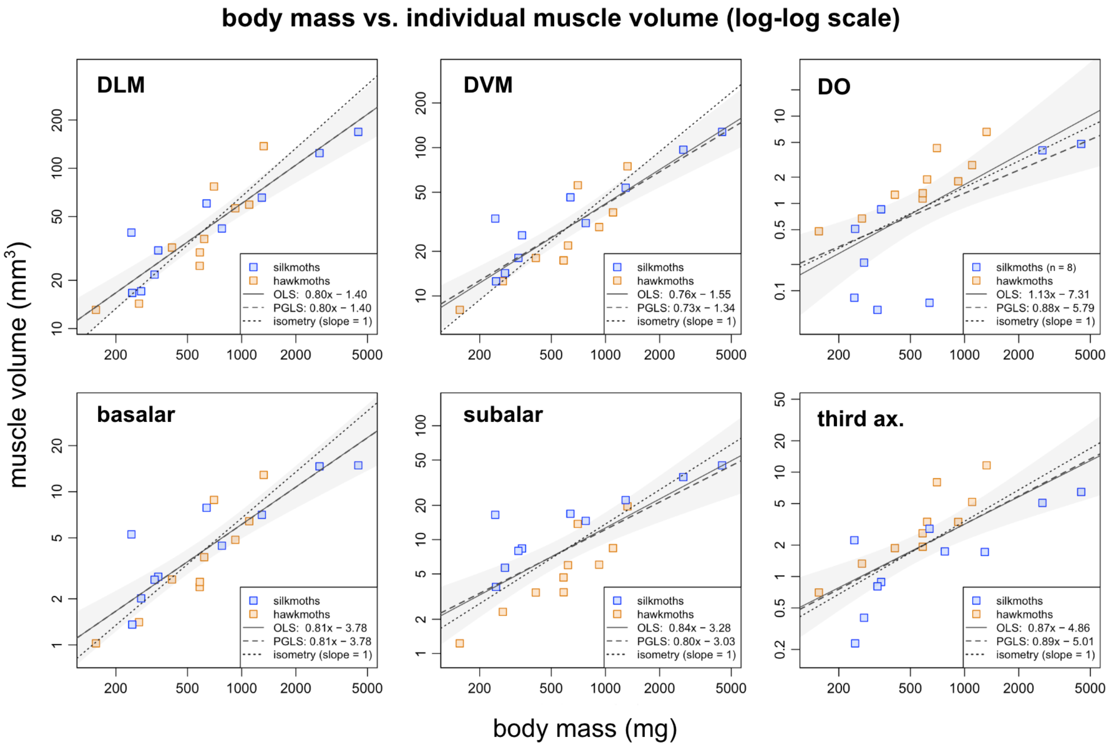


**Figure S1 Volume-body mass scaling for individual flight muscles.** Log-transformed body mass (mg) plotted against log-transformed volume ($mm^3$) of each flight muscle, pooled across both families ($n = 20$, except DO $n = 18$, as two species with entirely absent DOs were excluded from log-transformed analyses). Statistics are reported in Supp. Table S6. Solid lines indicate ordinary least-squares (OLS) regressions, shaded regions indicate OLS 95% confidence intervals, dashed lines indicate phylogenetic generalized least-squares (PGLS) regressions, and dotted lines indicate the isometric expectation (slope = 1). F-tests comparing observed PGLS slopes to isometry revealed significant hypoallometric scaling in the dorsolongitudinal muscle (DLM; slope = 0.80, $p = 0.035$) and dorsoventral muscle (DVM; slope = 0.73, $p = 0.006$). All other muscles exhibited slopes below 1, with the exception of the dorsal oblique (DO) under OLS, but did not differ significantly from isometry after phylogenetic correction. These results suggest that the hypoallometric scaling observed for total flight muscle volume is driven primarily by the two indirect flight power muscles.

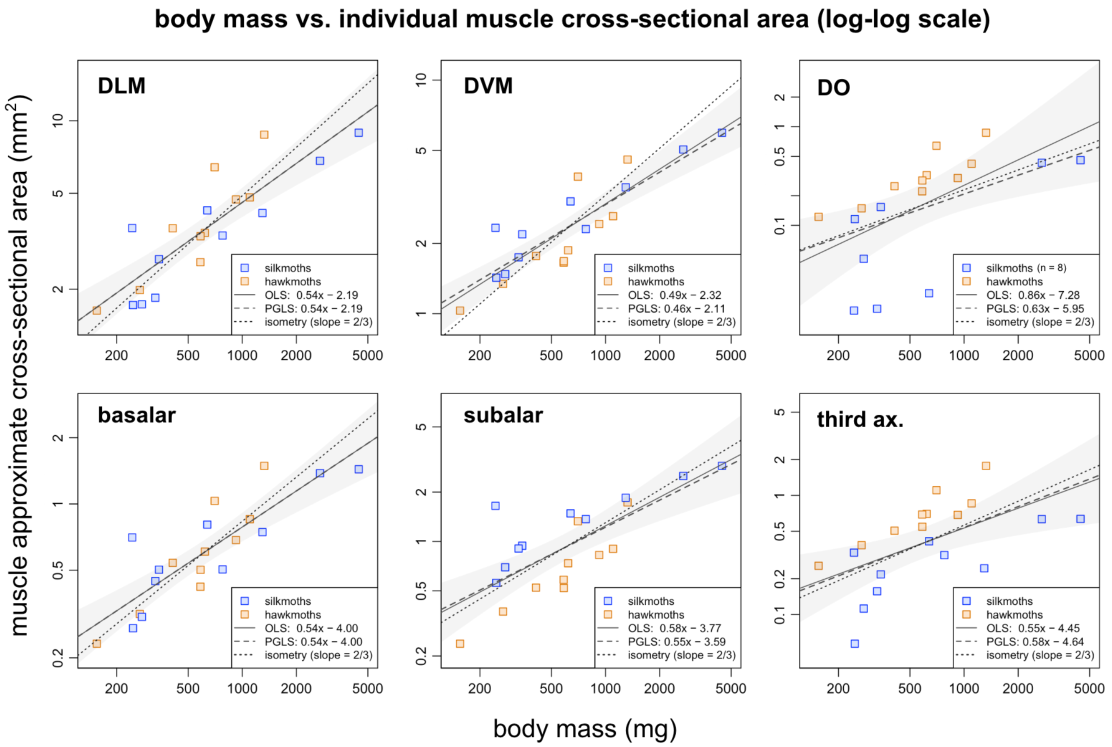


**Figure S2 Area-body mass scaling for individual flight muscles.** Log-transformed body mass (mg) plotted against approximate cross-sectional area ($mm^2$) for each flight muscle, pooled across both families [statistics in Supp. Table S6]. Solid lines indicate OLS regressions, shaded regions indicate OLS 95% confidence intervals, dashed lines indicate PGLS regressions, and dotted lines indicate the isometric expectation for area relative to body mass (slope = 2/3). Only the dorsoventral muscle (DVM) exhibited significant hypoallometric scaling relative to the isometric expectation after phylogenetic correction.

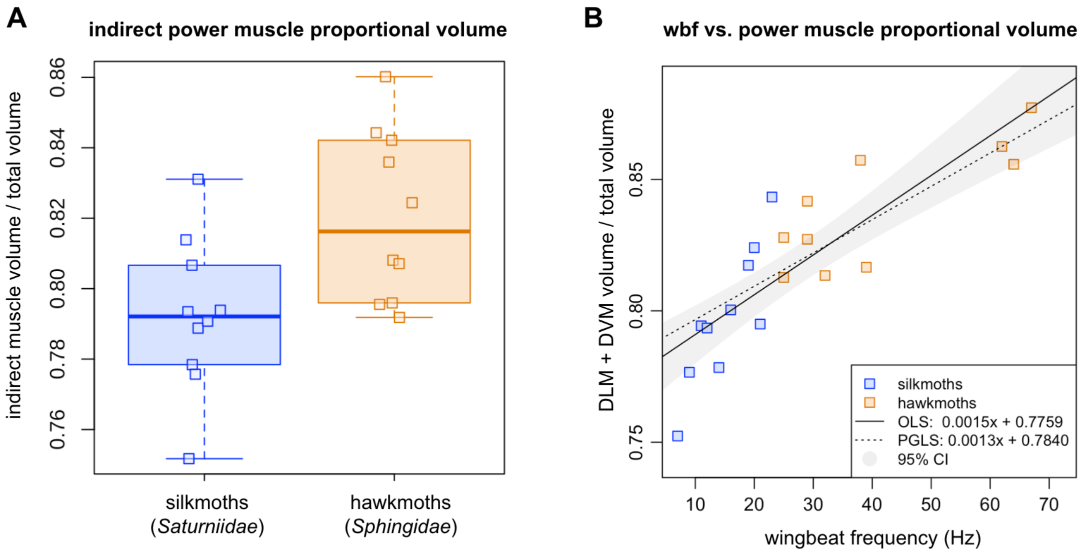


**Figure S3 Similar results persist when analyzing the indirect flight power muscles only** (DLM and DVM but not DO). The dorsal oblique (DO) is excluded because, although anatomically indirect, it is not generally considered a major power-producing flight muscle. We therefore repeated our analyses using only DLM and DVM volumes to verify that inclusion of the DO was not driving the observed relationships. **A.** Hawkmoths tend to have larger proportional volumes of indirect power muscle than silkmoths, consistent with the pattern observed in Fig. 3. This difference is significant under a conventional Welch two-sample t-test ($t$ = 2.73, $p = 0.014$), although a phylogenetic ANOVA is not significant ($p = 0.38$), matching the result obtained for total indirect muscle proportion. **B.** Wingbeat frequency remains positively associated with indirect power muscle proportion, indicating that the relationship in Fig. 3B is not driven by hawkmoths' larger DO muscles. The pooled relationship remains significant under OLS (slope = 0.0016, $p < 0.001$) as well as PGLS (slope = 0.0012, $p < 0.001$; $\lambda = 0.76$). Family-specific relationships remain significant for both hawkmoths (PGLS slope = 0.0011, $p = 0.005$) and silkmoths (PGLS slope = 0.0038, $p = 0.005$). Furthermore, the slopes of this relationship remain significantly different between the families under ANCOVA and phylogenetic ANCOVA (interaction $p = 0.005$). Based on these findings, we omit further analyses of DLM + DVM only.

| Family | Moth | Abbrev. | Body Mass (mg) | Wing Area (mm²) | Calculated WBF (Hz) | Measured WBF (Hz) I | Measured WBF (Hz) II |
|---|---|---|---|---|---|---|---|
| *Saturniidae* | *Actias luna* | AL | 776 | 1380 | 13.56 | 14 | 13.29 |
| | *Antheraea polyphemus* | AP | 638 | 1683 | 10.73 | 10 | 11.63 |
| | *Automeris io* | AI | 343 | 574 | 20.39 | 22 | 22.98 |
| | *Callosamia angulifera* | CA | 328 | 1586 | 8.82 | – | – |
| | *Citheronia regalis* | CR | 2707 | 1966 | 16.28 | 15 | – |
| | *Eacles imperialis* | EI | 4446 | 2080 | 18.80 | – | 15.90 |
| | *Hemileuca maia* | HM | 246 | 415 | 23.45 | – | – |
| | *Hyalophora euryalus* | HE | 1295 | 1948 | 12.41 | (13) | 12.99 |
| | *Samia cynthia* | SC | 244 | 1995 | 6.60 | – | – |
| | *Saturnia walterorum* | SW | 276 | 492 | 21.31 | – | – |
| *Sphingidae* | *Amorpha juglandis* | AJ | 269 | 339 | 28.65 | – | – |
| | *Amphion floridensis* | AF | 410 | 163 | 62.42 | – | 67.00 |
| | *Eumorpha achemon* | EA | 1101 | 638 | 29.23 | – | 33.60 |
| | *Hemaris thysbe* | HT | 585 | 188 | 63.80 | (62) | – |
| | *Hyles lineata* | HL | 922 | 433 | 37.71 | 35 | 38.97 |
| | *Manduca sexta* | MS | 1325 | 839 | 25.00 | 25 | – |
| | *Paonias myops* | PM | 619 | 341 | 39.41 | – | 41.88 |
| | *Proserpinus terlooi* | PT | 155 | 95 | 67.07 | – | – |
| | *Smerinthus ophthalmica* | SO | 584 | 420 | 32.37 | (33) | 34.88 |
| | *Sphinx chersis* | SpC | 701 | 619 | 25.17 | 31 | – |

**Table S1 Names and relevant metrics for twenty species** within the model clade *Bombycoidea*. Wing area data comes from Aiello, Tan, et al. (2021). Calculated wingbeat frequency, used in analyses, was obtained from body mass and wing area (see main text methods). Calculated frequencies generally correspond to prior free-flight frequency measurements taken by (I) Wold et al. (2024) and (II) Aiello, Sikandar, et al. (2021). Values in parentheses indicate a measurement from a different species of the same genus.

| | **DLM** | **DVM** | **DO** | **basalar** | **subalar** | **third ax.** | **total** |
|---|---|---|---|---|---|---|---|
| AL | 42.09 | 30.98 | 0.00 | 4.44 | 14.62 | 1.74 | 93.86 |
| AP | 60.32 | 46.21 | 0.07 | 7.86 | 16.86 | 2.87 | 134.19 |
| AI | 30.74 | 25.64 | 0.86 | 2.79 | 8.36 | 0.89 | 69.27 |
| CA | 21.72 | 18.04 | 0.06 | 2.66 | 7.97 | 0.80 | 51.26 |
| CR | 124.30 | 96.81 | 4.05 | 14.65 | 35.43 | 5.08 | 280.33 |
| EI | 168.61 | 127.68 | 4.80 | 14.87 | 44.89 | 6.46 | 367.30 |
| HM | 16.69 | 12.54 | 0.51 | 1.36 | 3.84 | 0.23 | 35.17 |
| HE | 65.58 | 53.64 | 0.00 | 7.07 | 22.25 | 1.72 | 150.26 |
| SC | 39.71 | 33.23 | 0.08 | 5.27 | 16.51 | 2.23 | 97.03 |
| SW | 17.09 | 14.25 | 0.21 | 2.01 | 5.67 | 0.40 | 39.63 |
| AJ | 14.31 | 12.58 | 0.67 | 1.41 | 2.32 | 1.33 | 32.62 |
| AF | 32.04 | 18.03 | 1.26 | 2.67 | 3.43 | 1.87 | 59.31 |
| EA | 59.30 | 36.58 | 2.75 | 6.42 | 8.42 | 5.18 | 118.64 |
| HT | 29.87 | 17.34 | 1.31 | 2.58 | 3.45 | 1.93 | 56.49 |
| HL | 56.28 | 29.12 | 1.79 | 4.85 | 6.02 | 3.34 | 101.41 |
| MS | 137.34 | 74.81 | 6.60 | 12.88 | 19.60 | 11.62 | 262.86 |
| PM | 36.22 | 21.88 | 1.88 | 3.74 | 5.97 | 3.34 | 73.04 |
| PT | 13.08 | 8.05 | 0.48 | 1.02 | 1.23 | 0.70 | 24.57 |
| SO | 24.63 | 17.39 | 1.14 | 2.38 | 4.66 | 2.59 | 52.79 |
| SpC | 76.95 | 55.77 | 4.30 | 8.84 | 13.74 | 8.01 | 167.61 |

**Table S2 Volume in mm$^3$ of six flight muscle pairs across twenty species,** along with total measured muscle volume per species. Each value is the sum volume of the two bilaterally symmetric muscles. The degree of asymmetry between bilateral muscle pairs is reported in Table S5.

| | DLM | DVM | DO | indirect | basalar | subalar | third ax. | direct |
|---|---|---|---|---|---|---|---|---|
| AL | 44.84 | 33.01 | 0.00 | 77.84 | 4.73 | 15.57 | 1.85 | 22.16 |
| AP | 44.95 | 34.44 | 0.05 | 79.44 | 5.85 | 12.57 | 2.14 | 20.56 |
| AI | 44.37 | 37.01 | 1.24 | 82.62 | 4.02 | 12.07 | 1.28 | 17.38 |
| CA | 42.38 | 35.20 | 0.12 | 77.69 | 5.20 | 15.54 | 1.57 | 22.31 |
| CR | 44.34 | 34.54 | 1.45 | 80.32 | 5.22 | 12.64 | 1.81 | 19.68 |
| EI | 45.90 | 34.76 | 1.31 | 81.97 | 4.05 | 12.22 | 1.76 | 18.03 |
| HM | 47.45 | 35.66 | 1.45 | 84.56 | 3.86 | 10.93 | 0.65 | 15.44 |
| HE | 43.65 | 35.70 | 0.00 | 79.35 | 4.70 | 14.81 | 1.14 | 20.65 |
| SC | 40.93 | 34.24 | 0.09 | 75.26 | 5.43 | 17.02 | 2.30 | 24.74 |
| SW | 43.13 | 35.94 | 0.53 | 79.61 | 5.08 | 14.30 | 1.02 | 20.39 |
| ***Sat.* mean** | **44.19** | **35.05** | **0.62** | **79.87** | **4.82** | **13.77** | **1.55** | **20.13** |
| AJ | 43.88 | 38.56 | 2.06 | 84.50 | 4.31 | 7.12 | 4.07 | 15.50 |
| AF | 54.03 | 30.40 | 2.12 | 86.55 | 4.51 | 5.79 | 3.16 | 13.45 |
| EA | 49.98 | 30.83 | 2.32 | 83.12 | 5.41 | 7.10 | 4.37 | 16.88 |
| HT | 52.89 | 30.71 | 2.32 | 85.91 | 4.56 | 6.11 | 3.42 | 14.09 |
| HL | 55.50 | 28.71 | 1.77 | 85.98 | 4.79 | 5.94 | 3.29 | 14.02 |
| MS | 52.25 | 28.46 | 2.51 | 83.22 | 4.90 | 7.46 | 4.42 | 16.78 |
| PM | 49.60 | 29.96 | 2.58 | 82.13 | 5.12 | 8.17 | 4.58 | 17.87 |
| PT | 53.24 | 32.78 | 1.95 | 87.97 | 4.17 | 5.00 | 2.86 | 12.03 |
| SO | 46.65 | 32.95 | 2.15 | 81.75 | 4.52 | 8.82 | 4.91 | 18.25 |
| SpC | 45.91 | 33.27 | 2.57 | 81.75 | 5.28 | 8.20 | 4.78 | 18.25 |
| ***Sph.* mean** | **50.39** | **31.66** | **2.23** | **84.29** | **4.76** | **6.97** | **3.99** | **15.71** |
| t-test | *** | ** | *** | *** | n.s. | *** | *** | *** |

**Table S3 Proportional volumes (%) of flight muscles by species,** with family-level means and t-test comparisons. Muscle proportions are calculated as bilateral muscle volume divided by the species' total measured muscle volume. Indirect = DLM + DVM + DO; direct = basalar + subalar + third axillary; direct = 100% - indirect. Two-tailed t-tests comparing silkmoth and hawkmoth proportions: *** $p < 0.001$, ** $p < 0.01$, * $p < 0.05$, n.s. = not significant.

| | DLM | DVM | DO | basalar | subalar | third ax. |
|---|---|---|---|---|---|---|
| AL | 6.30 | 6.82 | 0.00 | 4.40 | 5.34 | 2.76 |
| AP | 7.10 | 7.83 | 1.78 | 4.88 | 5.68 | 3.48 |
| AI | 5.77 | 5.99 | 2.79 | 2.77 | 4.44 | 2.04 |
| CA | 5.89 | 5.31 | 2.12 | 2.98 | 4.40 | 2.57 |
| CR | 9.12 | 9.73 | 4.70 | 5.31 | 7.06 | 4.02 |
| EI | 9.45 | 10.77 | 5.24 | 5.17 | 7.76 | 5.09 |
| HM | 4.85 | 4.43 | 2.21 | 2.49 | 3.44 | 2.01 |
| HE | 7.92 | 7.96 | 0.00 | 4.74 | 6.02 | 3.51 |
| SC | 5.54 | 7.31 | 3.04 | 3.74 | 5.01 | 3.37 |
| SW | 4.94 | 4.90 | 2.29 | 3.28 | 4.07 | 1.79 |
| AJ | 3.61 | 4.79 | 2.26 | 2.22 | 3.11 | 1.74 |
| AF | 4.47 | 5.16 | 2.52 | 2.48 | 3.29 | 1.85 |
| EA | 6.17 | 7.08 | 3.26 | 3.76 | 4.67 | 3.02 |
| HT | 4.50 | 5.21 | 2.30 | 2.57 | 3.32 | 1.77 |
| HL | 5.97 | 6.04 | 2.97 | 3.54 | 3.64 | 2.43 |
| MS | 7.85 | 8.25 | 3.80 | 4.32 | 5.66 | 3.28 |
| PM | 5.29 | 5.96 | 2.92 | 3.08 | 4.05 | 2.39 |
| PT | 4.01 | 3.96 | 1.97 | 2.21 | 2.58 | 1.37 |
| SO | 4.76 | 5.43 | 2.57 | 2.83 | 3.99 | 1.88 |
| SpC | 6.00 | 7.33 | 3.36 | 4.28 | 5.17 | 3.62 |

**Table S4 Average longitudinal-axis lengths in mm** of six flight muscles in twenty species.

| | DLM | DVM | DO | basalar | subalar | third ax. |
|---|---|---|---|---|---|---|
| AL | 3.34 | 3.08 | 0.00 | 0.51 | 1.37 | 0.32 |
| AP | 4.25 | 3.85 | 0.02 | 0.81 | 1.48 | 0.41 |
| AI | 2.66 | 2.57 | 0.15 | 0.50 | 0.94 | 0.22 |
| CA | 1.84 | 2.05 | 0.01 | 0.45 | 0.91 | 0.16 |
| CR | 6.82 | 6.39 | 0.43 | 1.38 | 2.51 | 0.63 |
| EI | 8.92 | 7.83 | 0.46 | 1.44 | 2.89 | 0.64 |
| HM | 1.72 | 1.88 | 0.12 | 0.27 | 0.56 | 0.06 |
| HE | 4.14 | 4.12 | 0.00 | 0.75 | 1.85 | 0.25 |
| SC | 3.59 | 2.72 | 0.01 | 0.71 | 1.65 | 0.33 |
| SW | 1.73 | 1.74 | 0.05 | 0.31 | 0.70 | 0.11 |
| AJ | 1.98 | 1.50 | 0.15 | 0.32 | 0.37 | 0.38 |
| AF | 3.58 | 3.10 | 0.25 | 0.54 | 0.52 | 0.51 |
| EA | 4.81 | 4.19 | 0.42 | 0.85 | 0.90 | 0.86 |
| HT | 3.32 | 2.87 | 0.29 | 0.50 | 0.52 | 0.55 |
| HL | 4.72 | 4.66 | 0.30 | 0.69 | 0.83 | 0.69 |
| MS | 8.75 | 8.32 | 0.87 | 1.49 | 1.73 | 1.77 |
| PM | 3.43 | 3.04 | 0.32 | 0.61 | 0.74 | 0.70 |
| PT | 1.63 | 1.65 | 0.12 | 0.23 | 0.24 | 0.26 |
| SO | 2.59 | 2.27 | 0.22 | 0.42 | 0.58 | 0.69 |
| SpC | 6.41 | 5.25 | 0.64 | 1.03 | 1.33 | 1.11 |

**Table S5 Approximate cross-sectional areas in $mm^2$** of six flight muscles in twenty species.

| trait | muscle | OLS slope | OLS $p_{reg}$ | OLS $F_{iso}$ | OLS $p_{iso}$ | PGLS slope | PGLS $p_{reg}$ | PGLS $F_{iso}$ | PGLS $p_{iso}$ |
|---|---|---|---|---|---|---|---|---|---|
| volume | DLM | 0.796 | *** | 5.23 | * | 0.796 | *** | 5.23 | * |
| | DVM | 0.765 | *** | 5.95 | * | 0.733 | *** | 9.67 | ** |
| | DO | 1.131 | *** | 0.18 | n.s. | 0.878 | *** | 0.42 | n.s. |
| | basalar | 0.810 | *** | 2.88 | n.s. | 0.810 | *** | 2.95 | n.s. |
| | subalar | 0.843 | *** | 0.92 | n.s. | 0.801 | *** | 3.67 | n.s. |
| | third ax. | 0.871 | *** | 0.48 | n.s. | 0.893 | *** | 0.47 | n.s. |
| area | DLM | 0.537 | *** | 3.09 | n.s. | 0.537 | *** | 3.09 | n.s. |
| | DVM | 0.493 | *** | 7.65 | * | 0.461 | *** | 16.64 | *** |
| | DO | 0.857 | ** | 0.42 | n.s. | 0.634 | ** | 0.04 | n.s. |
| | basalar | 0.545 | *** | 2.38 | n.s. | 0.545 | *** | 2.38 | n.s. |
| | subalar | 0.578 | *** | 0.57 | n.s. | 0.549 | *** | 2.22 | n.s. |
| | third ax. | 0.553 | ** | 0.38 | n.s. | 0.581 | *** | 0.44 | n.s. |

**Table S6 Scaling of body mass with volume and area** of individual flight muscles. Regression significance ($p_{reg}$) tests whether slope differs from zero. Isometry tests compare observed slopes with the isometric expectation (volume isometric slope = 1, area isometric slope = 2/3), with F-statistic ($F_{iso}$) and significance of slope difference ($p_{iso}$).

| | | **DLM** | **DVM** | **DO** | **indirect** | **basalar** | **subalar** | **third ax.** | **direct** |
|---|---|---|---|---|---|---|---|---|---|
| *Saturniidae* | median | 44.37 | 35.02 | 0.32 | 79.54 | 4.91 | 13.47 | 1.67 | 20.46 |
| | Q1 | 43.28 | 34.54 | 0.06 | 78.16 | 4.21 | 12.30 | 1.18 | 18.39 |
| | Q3 | 44.87 | 35.66 | 1.29 | 81.62 | 5.22 | 15.43 | 1.84 | 21.84 |
| | IQR | 1.66 | 1.23 | 1.23 | 3.34 | 1.01 | 3.05 | 0.67 | 3.34 |
| *Sphingidae* | median | 51.14 | 30.84 | 2.23 | 83.93 | 4.68 | 7.11 | 4.22 | 16.07 |
| | Q1 | 47.42 | 30.14 | 2.07 | 82.37 | 4.51 | 5.98 | 3.32 | 14.02 |
| | Q3 | 53.17 | 32.87 | 2.46 | 86.00 | 5.06 | 7.99 | 4.54 | 17.63 |
| | IQR | 5.77 | 2.84 | 0.39 | 3.59 | 0.55 | 2.01 | 1.22 | 3.59 |

**Table S7 Boxplot summary statistics for flight muscle proportional volumes (%),** by family. Median, first (Q1) and third (Q3) quartiles, and interquartile range (IQR) for each muscle. Values are bilateral muscle volume as a percentage of total measured muscle volume ($n$ = 10 per family). Indirect = DLM + DVM + DO; direct = basalar + subalar + third axillary.

| | | DLM | DVM | DO | basalar | subalar | third ax. |
|---|---|---|---|---|---|---|---|
| | *Sat.* mean | 1.41 | 1.48 | 0.51 | 0.84 | 1.12 | 0.63 |
| **n.d. length** | *Sph.* mean | 1.22 | 1.38 | 0.65 | 0.72 | 0.92 | 0.53 |
| | t-test p-val. | *** | ** | n.s. | ** | *** | ** |
| | *Sat.* mean | 0.16 | 0.15 | 0.005 | 0.029 | 0.061 | 0.012 |
| **n.d. area** | *Sph.* mean | 0.21 | 0.18 | 0.017 | 0.033 | 0.038 | 0.037 |
| | t-test p-val. | *** | *** | *** | ** | *** | *** |

**Table S8 T-test comparisons of non-dimensional length and cross-sectional area.** Each metric is normalized to an exponent of total flight muscle volume to control for body size scaling effects: n.d. length = length (mm) $\div$ total volume$^{1/3}$ (mm), and n.d. area = area (mm$^2$) $\div$ total volume$^{2/3}$ (mm$^2$). Silkmoths have higher normalized lengths for each muscle except the DO, which does not exhibit a significant difference in length. Hawkmoths have higher normalized areas for each muscle except the subalar, which is larger in silkmoths. Two-tailed t-test: *** $p < 0.001$, ** $p < 0.01$, * $p < 0.05$.

### References


Aiello, B. R., Sikandar, U. B., Minoguchi, H., Bhinderwala, B., Hamilton, C. A., Kawahara, A. Y., & Sponberg, S. (2021). The evolution of two distinct strategies of moth flight. *Journal of The Royal Society Interface*, *18*(185), 20210632.

Aiello, B. R., Tan, M., Bin Sikandar, U., Alvey, A. J., Bhinderwala, B., Kimball, K. C., Barber, J. R., Hamilton, C. A., Kawahara, A. Y., & Sponberg, S. (2021). Adaptive shifts underlie the divergence in wing morphology in bombycoid moths. *Proceedings of the Royal Society B: Biological Sciences*, *288*(1956), 20210677.

Bartholomew, G. A., & Casey, T. M. (1978). Oxygen Consumption of Moths During Rest, Pre-Flight Warm-Up, and Flight in Relation to Body Size and Wing Morphology. *Journal of Experimental Biology*, *76*(1), 11–25.

Beenakkers, A., Van Der Horst, D., & Van Marrewijk, W. (1984). Insect flight muscle metabolism. *Insect Biochemistry*, *14*(3), 243–260.

Casey, T. M. (1981). A Comparison of Mechanical and Energetic Estimates of Flight Cost for Hovering Sphinx Moths. *Journal of Experimental Biology*, *91*(1), 117–129.

Dudley, R. (2000). *The Biomechanics of Insect Flight: Form, Function, Evolution*. Princeton University Press.

Ellington, C. P. (1985). Power and efficiency of insect flight muscle. *Journal of Experimental Biology*, *115*(1), 293–304.

Fedorov, A., Beichel, R., Kalpathy-Cramer, J., Finet, J., Fillion-Robin, J.-C., Pujol, S., Bauer, C., Jennings, D., Fennessy, F., Sonka, M., Buatti, J., Aylward, S., Miller, J. V., Pieper, S., & Kikinis, R. (2012). 3D Slicer as an image computing platform for the Quantitative Imaging Network. *Magnetic Resonance Imaging*, *30*(9), 1323–1341.

Harrison, J. F. (2018). Approaches for testing hypotheses for the hypometric scaling of aerobic metabolic rate in animals. *American Journal of Physiology-Regulatory, Integrative and Comparative Physiology*, *315*(5), R879–R894.

Harrison, J. F., Biewener, A., Bernhardt, J. R., Burger, J. R., Brown, J. H., Coto, Z. N., Duell, M. E., Lynch, M., Moffett, E. R., Norin, T., Pettersen, A. K., Smith, F. A., Somjee, U., Traniello, J. F. A., & Williams, T. M. (2022). White Paper: An Integrated Perspective on the Causes of Hypometric Metabolic Scaling in Animals. *Integrative and Comparative Biology*, *62*(5), 1395–1418.

Hulbert, A. (2014). A Sceptics View: “Kleiber’s Law” or the “3/4 Rule” is neither a Law nor a Rule but Rather an Empirical Approximation. *Systems*, *2*(2), 186–202.

Kleiber, M. (1932). Body size and metabolism. *Hilgardia*, *6*(11), 315–353.

Lasso, A. (2025, June). Lassoan/SlicerSegmentEditorExtraEffects [original-date: 2017-03-21T15:33:53Z].

Niven, J. E., & Scharlemann, J. P. (2005). Do insect metabolic rates at rest and during flight scale with body mass? *Biology Letters*, *1*(3), 346–349.

Stern, D. L., & Emlen, D. J. (1999). The developmental basis for allometry in insects. *Development*, *126*(6), 1091–1101.

Suarez, R. K. (2000). Energy Metabolism during Insect Flight: Biochemical Design and Physiological Performance. *Physiological and Biochemical Zoology*, *73*(6), 765–771.

Tolfrey, K., Barker, A., Thom, J. M., Morse, C. I., Narici, M. V., & Batterham, A. M. (2006). Scaling of maximal oxygen uptake by lower leg muscle volume in boys and men. *Journal of Applied Physiology*, *100*(6), 1851–1856.

Weibel, E. R. (2002). The pitfalls of power laws. *Nature*, *417*(6885), 131–132.

White, C. R., Cassey, P., & Blackburn, T. M. (2007). Allometric exponents do not support a universal metabolic allometry. *Ecology*, *88*(2), 315–323.

Wold, E. S., Aiello, B., Harris, M., Bin Sikandar, U., Lynch, J., Gravish, N., & Sponberg, S. (2024). Moth resonant mechanics are tuned to wingbeat frequency and energetic demands. *Proceedings of the Royal Society B: Biological Sciences*, *291*(2025), 20240317.
